# Pathogenicity effector candidates and accessory genome revealed by pan-genomic analysis of *Parastagonospora nodorum*

**DOI:** 10.1101/2021.09.01.458590

**Authors:** Darcy A. B. Jones, Kasia Rybak, Stefania Bertazzoni, Kar-Chun Tan, Huyen T. T. Phan, James K. Hane

## Abstract

The wheat pathogen *Parastagonospora nodorum* has emerged as a model necrotrophic fungal species with growing genomic resources. Recent population-level pan-genome studies were leveraged to provide novel insights into pathogen evolution and effector-like gene contents relevant to local crop disease outbreaks. In this study, we examined 156 isolates representing a regional population from the Western Australian (WA) wheat-belt region, and 17 internationally sourced isolates. We observed a highly diverse local population, within which were numerous small and highly similar clusters of isolates from hotter and drier regions. Pan-genome assembly and orthologous gene datasets resulted in 3579 predicted effector candidates, 2291 of which exhibited presence-absence variation (PAV) across the population, and 1362 were specific to WA isolates. There was an abundance of mutations (including repeat-induced point mutation (RIP)), distributed in ‘hot-spots’ within the pan-genomic landscape that were rich in effector candidates. Three characterised effector loci (*ToxA*, *Tox1* and *Tox3*) were located within sub- telomeric regions of lower diversity, but were nestled within larger high-diversity regions. RIP was widespread across the genome, but non-synonymous RIP-like mutations were strongly selected against. These improved bioinformatic resources for *P. nodorum*, represent progressive advancements in fungal pan-genomics, with a view towards supporting region-specific surveillance of host-pathogen interactions.

## Introduction

*Parastagonospora nodorum* is a necrotrophic pathogen causing septoria nodorum blotch (SNB) of wheat (*Triticum spp.*) (Solomon et al. 2006) which leads to significant yield losses in Australia (Murray and Brennan 2009). *P. nodorum* is primarily spread by infected seed, infested debris or by wind dispersed sexual ascospores (Solomon et al. 2006). Secondary infections can occur when water splash spreads asexual pycnidiospores to higher leaves and glumes, causing further necrotic patches and crop loss. *P. nodorum* is observed to be highly diverse in the field (Stukenbrock et al. 2006, McDonald et al. 2012), and appears to regularly reproduce sexually (Murphy et al. 2000, Bathgate and Loughman 2001, Sommerhalder et al. 2006). This suggests a high adaptive capacity of *P. nodorum* populations, where application of strong selective pressures may be quickly overcome by extant diversity.

*P. nodorum* infection heavily relies on the activity of necrotrophic effector proteins (NEs), which are secreted into the host and cause cell death upon recognition by host susceptibility (S)- proteins (Tan et al. 2010). Three NEs have been characterised in *P. nodorum* to date: ToxA (Liu et al. 2006), Tox1 (Liu et al. 2012) and Tox3 (Liu et al. 2009). At least five additional host specific necrosis phenotypic interactions have been described in the *P. nodorum*-wheat pathosystem (Friesen et al. 2007, Friesen et al. 2008, Abeysekara et al. 2009, Zhang et al. 2011, Friesen et al. 2012, Gao et al. 2015, Shi et al. 2015, Phan et al. 2018), indicating the presence of additional undiscovered NEs. Identification of the *ToxA* NE has led to deployment of a ToxA resistant wheat cultivar (Tan et al. 2014), and the presence of additional major disease resistance quantitative trait loci (QTL) encourages further development of disease resistant cultivars. However, the known epistatic interactions of Tox1 and Tox2 over Tox3 (Friesen et al. 2007, Phan et al. 2016) indicates that the contributions of different effector-receptor interactions to virulence are complex, and that reliable markers of S-genes and knowledge of the epistasis interactions are important for future disease breeding efforts. The discovery of novel NEs in *P. nodorum* (and other fungal pathogens) remains an important element of crop protection research (Vleeshouwers and Oliver 2014). Fungal effector discovery relies heavily on genomic and bioinformatic resources (Jones et al. 2018), and the increased accessibility of sequencing has resulted in a considerable increase in the rate of effector discovery (Kanja and Hammond-Kosack 2020).

*P. nodorum* was among the first fungal species for which a reference genome sequence was generated (for the Western Australian (WA) isolate SN15) (Hane et al. 2007), and was the first species within the class Dothideomycetes, which comprises several prominent cereal necrotroph and hemibiotroph species (Ohm et al. 2012, Aylward et al. 2017). The SN15 reference isolate has predominated molecular plant pathology studies of *P. nodorum* since, becoming an important model for the cereal necrotrophs (Solomon et al. 2006). Significant resources have accumulated over time for the SN15 isolate and *P. nodorum* in general, including transcriptomic (Hane et al. 2007, Ipcho et al. 2012, Syme et al. 2016, Richards et al. 2018, Jones et al. 2019), proteomic (Bringans et al. 2009, Syme et al. 2016), and metabolomic (Lowe et al. 2008, Gummer et al. 2013, Chooi et al. 2014, Muria-Gonzalez et al. 2020) datasets. Notable recent additions to the growing pool of *P. nodorum* data include the chromosome-scale genome assemblies of SN15 and 3 other reference isolates (Richards et al. 2018, Bertazzoni et al. 2021).

### Parastagonospora nodorum pan-genomics

Improving cost and availability of genome and transcriptome sequencing has significantly advanced plant pathology, enabling conceptual shifts from the focused study of a single reference isolate, to large-scale comparative genomics between numerous isolates over a few short years. Three pan-genomic comparative studies of *P. nodorum* have been conducted to date, comparing global collections of isolates (Syme et al. 2018, Pereira et al. 2020, Bertazzoni et al. 2021), and populations within the USA (Richards et al. 2019). Syme *et al*. (Syme et al. 2018) compared the genomes of *P. nodorum* isolates from Iran, Finland, Sweden, Switzerland, South Africa, the USA, and Australia. They observed frequent presence-absence variation in the effectors *ToxA*, *Tox1*, and *Tox3*, and possible accessory genomic regions with potential roles in virulence. Additionally, numerous genes were observed to be under positive selection pressure, including several effector candidate proteins. They observed distinct regions with large numbers of mutations and some regions with consistently high dN/dS ratios, indicating positive selection and enrichment of adaptively relevant genes in specific genomic regions. Pereira *et al*. (Pereira et al. 2020) sequenced the genomes of *P. nodorum* isolates from Australia, Iran, South Africa, Switzerland, and the USA to investigate *P. nodorum* adaptation to fungicide use. The highest genetic diversity was found in the Iranian population, consistent with the hypothesis that *P. nodorum* co-evolved with wheat during early domestication (Ghaderi et al. 2020), while low diversity was observed within the Australian isolates. A genome-wide association study identified several loci correlated with azole resistance, with higher incidence of fungicide resistance and the identified resistance associated alleles in the Swiss population. Richards *et al*. (Richards et al. 2019) sequenced 197 *P. nodorum* isolates collected from Spring, Winter and Durum wheat cultivars across the growing region of the USA. Two major US sub-populations were identified corresponding to geographical features and wheat lines grown, with one sub- population almost completely lacking *ToxA*. Both populations had diversified at different loci, indicating distinct selective pressures and resulting in different sets of effector candidates predicted by genome-wide association. The USA population study also highlighted several patterns of gene presence-absence variation (PAV) between isolates. The known effector genes *ToxA, Tox1 and Tox3* were absent in 37%, 5% and 41% of US isolates respectively. Gene PAV was commonly associated with transposons in USA isolates, yet there was no significant association with frequency of secreted or effector-like proteins.

Collectively, these genomic studies have highlighted the high diversity of *P. nodorum* genomes, and that studies focusing only on reference isolates are missing considerable amounts of information. The 400 kb accessory chromosome 23 (AC23), which is missing in the avirulent isolate Sn79-1087, may have a role in host-specific virulence (Bertazzoni et al. 2018) and has a high background rate of mutation (Richards et al. 2018, Bertazzoni et al. 2021). Similarly, regional biases for high AT-base content and RIP-like activity suggests the presence of rapidly mutating accessory regions, primarily within repeat-rich stretches of AC23 and sub-telomeric regions (Richards et al. 2018, Syme et al. 2018, Bertazzoni et al. 2021). Numerous effector candidates have been derived from these genomic studies, utilising features such as signatures of positive selection across isolates, similarity to known effectors, signal peptides, EffectorP results (Sperschneider et al. 2016, Sperschneider et al. 2018), genomic context (e.g. AT-richness or distance to TEs), presence or absence in avirulent isolates, and genome wide association (Richards et al. 2018, Syme et al. 2018, Richards et al. 2019, Bertazzoni et al. 2021). In the Australian reference isolate SN15, many of these genomic features have been combined with a broad range of experimental and bioinformatic indicators to refine effector candidate lists, including: *in planta* gene expression (Jones et al. 2019), putative lateral gene transfer with other cereal-pathogenic fungi (https://effectordb.com), and predicted effector-like gene and protein properties (Syme et al. 2018, Bertazzoni et al. 2021).

### The Western Australian *P. nodorum* pan-genome

Despite the long history of study of the Australian *P. nodorum* reference isolate SN15, relatively little was known about the genomic diversity of the *P. nodorum* population in WA until recently. A study of 28 SSR loci compared a WA population of 155 isolates collected over 44 years, and contrasted this population with 23 international isolates sourced from France and the USA (Phan et al. 2020). This SSR study identified two core admixed clusters in WA, with three low-diversity satellite clusters that were geographically and temporally restricted. The population shift broadly correlated with historical shifts in wheat cultivar preference, particularly after the mass adoption of the ToxA insensitive cultivar “Mace” in 2013 which covers nearly 70% of the area sown (Trainor et al. 2018). Although wheat cultivar disease resistance has increased over time, more recently sampled isolates from emergent clusters were more pathogenic than older isolates.

In this study, we further dissect the evolutionary history of a WA *P. nodorum* population previously analysed by Phan *et al*. (Phan et al. 2020) using whole genome sequencing. Further, we compare these genomes with isolates previously sequenced by Syme *et al*. (Syme et al. 2018), and identify novel effector candidates in *P. nodorum*. Guided by phylogeographic and population structure analyses, we observe a highly diverse population in WA, with numerous small highly similar clusters collected from hotter and drier regions of Western Australia. We present the first report of novel sequences and genes within the growing pool of *P. nodorum* pangenome data, specific to locally-adapted isolates. We also combine new WA pangenome data with existing resources to define new effector candidate orthologous groups, extending effector candidate discovery beyond reference isolates. Overall, this growing wealth of new pathogen population data has enhanced our understanding of the pathogenicity gene content, genome architecture and population dynamics for the model cereal necrotroph, *P. nodorum*.

## Results

### Quality control of input sequence data

A total of 156 Western Australian *P. nodorum* isolates described previously (Syme et al. 2018, Phan et al. 2020) were sequenced using short-read Illumina sequencing. The majority of sequencing read pairs were assigned by Kraken2 to either *Parastagonospora nodorum* (average 74.1%) or were unclassified (average 25.6%) (table S2). A small number of read pairs matched other organisms across kingdoms, but no fastq files had more than 1% of reads assigned to any non-fungal taxon nor were any consistent trends observed. Sequencing read quality was generally high, with only a single lane of sequencing (of three) for the isolate WAC13068 failing QC because of low base quality (data S1). Mean insert sizes for isolates sequenced with 125 bp reads ranged between 441 and 709 bp (mean 632 bp), with standard deviations between 263bp and 760bp. Insert sizes for isolates sequenced with 150bp reads varied between 275bp and 321bp (mean 305 bp) with standard deviations between 208 bp and 295 bp.

### Prediction of mutations across the *P. nodorum* pan-genome relative to the SN15 reference isolate

Short variants (SNPs, insertions, and deletions) were predicted from aligned short-read sequenced isolates from this study and 15 previously published additional isolates (Syme et al. 2018) (table S1) using the SN15 genome as a reference (Bertazzoni et al. 2021), yielding 895,000 variant loci after filtering (table S3, data S2-S8) corresponding to 1 mutation for every 41 bases in the genome on average. The majority of variants were SNPs (830,761), compared to 33,943 and 30,296 insertions and deletions. Most SNPs were C↔T or G↔A mutations (296,688 and 296,927, respectively), with an overall transition to transversion ratio of 2.5. Relative to the genome of the SN15 reference isolate, 7.4% of variants were within exon features, 3236 mutations resulted in truncation (gain of stop), 660 mutations resulted in loss of a stop codon, 466 mutations resulting in loss of start codon, 249 and 310 mutations were in splice site acceptor or donor sites respectively, 9896 were frameshift variants, 148,914 were missense mutations, 205,602 were synonymous variants, and there were 1648 and 1591 disruptive in-frame deletions and insertions, respectively.

### Phylogeny and structure of the local Western Australian *P. nodorum* population

A subset of SNPs for phylogenetic and population genetics analyses were selected (maximum missing genotypes of 30%, minimum non-major allele frequency of 0.05, and filtering correlated loci within 10 Kb) resulting in 45,194 SNP loci from all illumina sequenced isolates.

A phylogenetic tree was estimated from these SNPs using IQTree (Minh et al. 2020) (Fig 1). The resulting tree grouped isolates from WA and non-Australian isolates into distinct clades. Leaf lengths were generally long within the WA clade, however, six groups of isolates with very short branch lengths were observed, with sizes ranging from 3 to 14 isolates (Data S9, Fig S1). Some internal tree nodes had low UFBoot support values (< 95), indicating that some of the high level relationships were poorly resolved. A major split with high SH-aLRT but low UFBoot support was observed, which tended to separate isolates with long branch lengths from a diverse range of locations from a second clade comprised mostly of several highly similar clades which were generally collected from northern regions of the WA wheat growing area (Geraldton and Dandaragan). No obvious correspondence between effector haplotype profiles and high level phylogenetic clades was observed; but at lower level clades effector haplotype profiles appear to be conserved, particularly where member isolates have short branch lengths (Fig 1).

**Fig 1.**
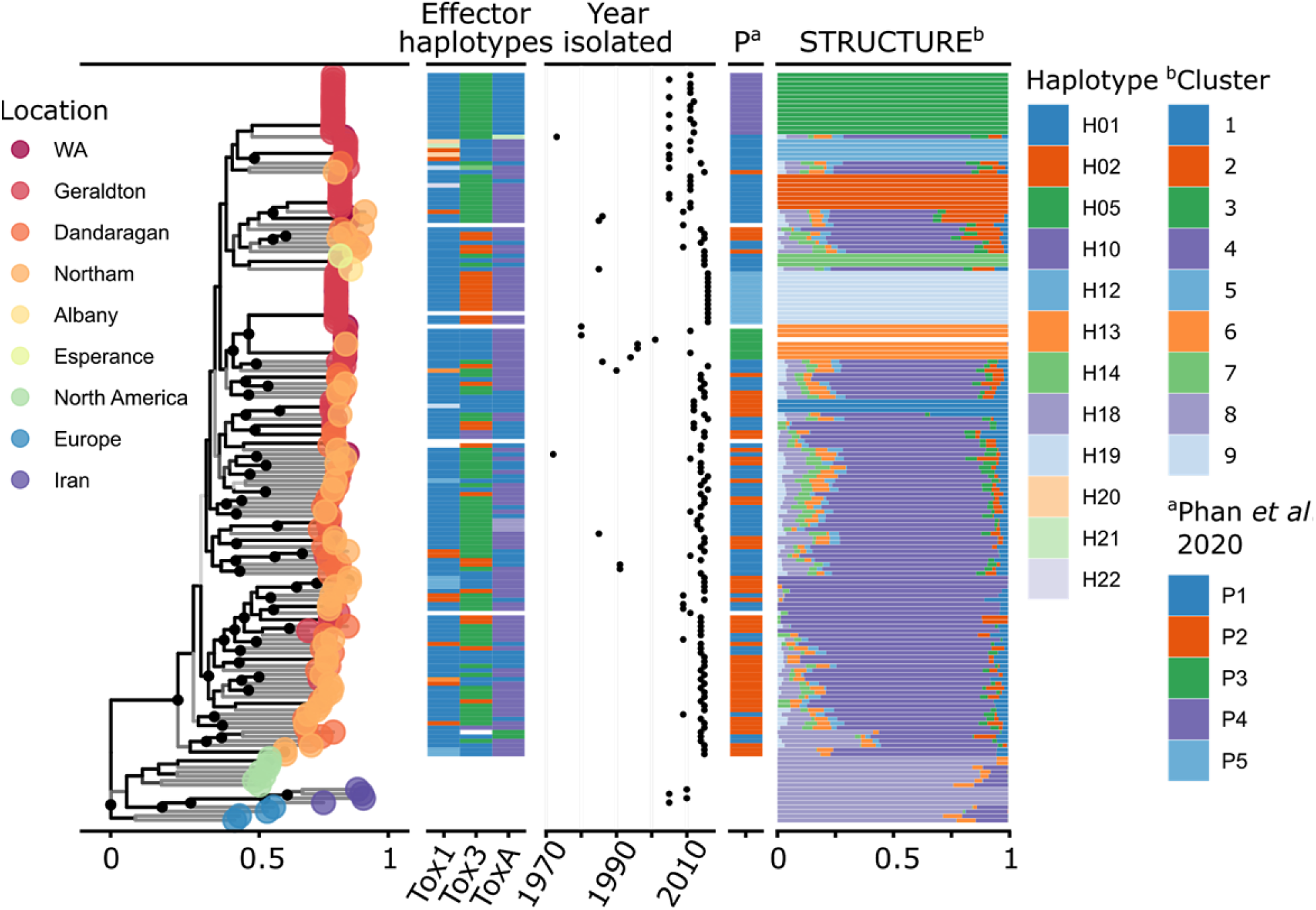
The structure and features of the Western Australian (WA) *Parastagonospora nodorum* population. The tree on the left shows the predicted phylogeny of WA and internationally- sampled *P. nodorum* isolates, with colours corresponding to sampling regions, summarised from Fig S1. Dots on the tree indicate where clades have >= 95% UFBoot confidence, and clade shade indicates SH-aLRT scores, with black indicating high support. Effector haplotype profiles for three confirmed effector loci are shown based on data from Phan *et al*. (Phan et al. 2020). A haplotype with white indicates that the isolate has not been haplotyped, or was unable to be amplified by PCR. On the right shows the results of population structure analysis, with colours indicating discrete population clusters, and the relative size of the bars indicating posterior probability of an isolate belonging to that cluster. The section indicated as “P” shows the clusters assigned to that isolate by the SSR study from Phan *et al*. (Phan et al. 2020), where white indicates isolates that were not present in that study

Analysis of population structure from SNP data with STRUCTURE (Fig 1, Table S4), predicted nine clusters, with the non-Australian isolates forming a single cluster (Structure cluster 8). Within the WA isolates, a single main population was observed (cluster 4) with 7 small satellite clusters. Few isolates were unambiguously assigned to the main cluster (21 of 102 had a posterior probability of > 0.85), with small but appreciable posterior probabilities contributed by satellite clusters and the international cluster. Noting this, we refer to isolates with the highest posterior probability of assignment to cluster 4 as members. These members generally had long branch lengths in the phylogenetic tree and were found in multiple clades. This main cluster corresponds to the two main clusters (1 and 2) presented in Phan et al. (Phan et al. 2020) (referred to hereafter as clusters P1-5) (Fig S2 and S3). The seven remaining clusters identified by STRUCTURE corresponded to clades in the phylogenetic tree where all leaves had very short lengths, indicating that all members were highly similar to each other. Cluster 1 consists of 3 isolates collected from Geraldton in 2012 (Fig S4 and S5). Cluster 2 consists of 8 isolates collected from Geraldton in 2005 and 2011. Cluster 3 consists of 14 isolates collected from Geraldton in 2005, 2011, and 2012. Cluster 5 consists of 5 isolates collected from Geraldton in 2005 and 2011. Cluster 6 consists of 7 isolates collected from Geraldton, South Perth, or WA (unknown specific location) between the years 1980 and 2011. Cluster 9 consists of 12 isolates collected from Mingenew (near Geraldton) or WA in 2016. Clusters 3, 6, and 9 correspond to the satellite clusters P3, P4, and P5 presented by Phan *et al*. (Phan et al. 2020), respectively.

The remaining clusters further separate the main populations identified by Phan *et al*. (Phan et al. 2020), with cluster 1 separating from P2, and clusters 7, 5, and 2 separating from P1. The Tox1 effector haplotypes of members in cluster 5 were highly variable and contained rare variants in the WA population.

Population diversity statistics indicated that individuals within clusters except the core WA cluster (4) and the international cluster (8) are highly similar (Table 1, Table S4). F_ST_ and average locus expected heterozygosity suggests that the alleles are nearly fixed in these clusters. The mean locus G^′^_ST_ (a normalised variant of F_ST_ that accounts for multi-allelic markers) (Hedrick 2005) was 0.44, indicating that there is some differentiation between subpopulations. To reduce the effects of nearly identical isolates on local diversity estimates, reduced multilocus genotypes (MLGs) were defined using the IQTree maximum likelihood (ML) distance estimates using complete linkage clustering and a cutoff threshold of 0.1 (Table 1). All clusters other than 4 and 8 (the main WA population and the international isolates) were composed of a single MLG using this strategy. Cluster 4 comprised 89 MLGs (of 102 individuals) and cluster 8 comprised 14 MLGs (of 15 individuals). For each sampling location and year, a single individual of each MLG was selected for further comparison, referred to as the “clone-corrected” subset. A mantel test comparing the ML distance matrix with the distance matrix derived from the isolate sampling GPS coordinates indicated no significant correlation between genetic distance and geographic distance in the WA “clone-corrected” population (Mantel test, 999 replications, p-value = 0.381). Principal components analysis (PCA) of the clone corrected samples showed a clear separation of international isolates from those of WA in PC1 which explained 4.7% of the total variance, but no other principal components showed structure in the data correlated with sampling location or year (Fig S6 and S7).

**Table 1.** Population diversity statistics based on 45,194 SNP variant loci (44,532 biallelic) from the *P. nodorum* population. Isolates are assigned to populations based on the STRUCTURE cluster with the highest posterior probability. No isolates had the same SNP profile, so multilocus genotypes (MLGs) were defined as having a genetic distance (estimated by IQTree) less than 0.1. The column “# loci” indicates the number of loci within a subpopulation that were variable and had no missing genotypes. The mean locus Simpson’s index (Mean λ. AKA expected heterozygosity) is calculated for every locus under analysis, for a biallelic SNP in a haploid organism the maximum possible value is 0.5. The Simpson’s index (λ), Stoddart- Taylor’s index (G), and Shannon’s diversity (H) are calculated based on MLGs. The index of association (IA) and it’s normalised form ( rd) indicate linkage disequilibrium in each population cluster, and was calculated only on variant loci with no missing data (# loci) within each cluster. rd p-values indicate the result of a permutation test (999 permutations) for higher rd than would be expected by random allele distribution across isolates.

| | # isolates | # MLG | # loci | $F_{ST}$ | Mean $\lambda$ | $\lambda$ | G | H | $I_A$ | $\bar{r}_d$ | $\bar{r}_d$ p-value |
| --- | --- | --- | --- | --- | --- | --- | --- | --- | --- | --- | --- |
| Cluster 1 | 3 | 1 | 155 | 0.99 | 0.002 | 0 | 1 | 0 | 0.11 | 7e-3 | 0.325 |
| Cluster 2 | 8 | 1 | 149 | 0.99 | 0.002 | 0 | 1 | 0 | 3.77 | 0.03 | 0.001 |
| Cluster 3 | 14 | 1 | 156 | 0.99 | 0.003 | 0 | 1 | 0 | 0.58 | 0.004 | 0.026 |
| Cluster 4 | 102 | 89 | 10353 | 0.05 | 0.311 | 0.99 | 81.28 | 4.45 | 1.53 | 0.006 | 0.001 |
| Cluster 5 | 5 | 1 | 114 | 0.99 | 0.002 | 0 | 1 | 0 | 2.38 | 0.021 | 0.001 |
| Cluster 6 | 7 | 1 | 33 | 1.0 | 0.001 | 0 | 1 | 0 | 1.27 | 0.040 | 0.021 |
| Cluster 7 | 3 | 1 | 114 | 0.99 | 0.001 | 0 | 1 | 0 | -0.03 | -2e-3 | 0.363 |
| Cluster 8 | 15 | 14 | 6695 | 0.0 | 0.314 | 0.92 | 13.24 | 2.62 | 11.85 | 0.013 | 0.001 |
| Cluster 9 | 12 | 1 | 87 | 1.0 | 0.001 | 0 | 1 | 0 | 0.54 | 0.006 | 0.048 |
| Total | 169 | 110 | 45194 |  | 0.320 |  |  |  |  |  |  |

Permutation tests of the *r_d_* index of association indicated that all clusters except 1 and 7 were in linkage disequilibrium (999 replications, p-value < 0.05) (Table 1). Clusters 1 and 7 both contain only three members, so the test may be underpowered in those cases. Repeated tests for clusters 4 and 8 using the “clone corrected” subset were also significant, indicating that linkage disequilibrium was not associated with isolate clonality.

### Comparative genomics across the local Western Australian *P. nodorum* population indicated telomeric or transposon-rich mutation ‘hotspots’

Short variant mutation frequencies were observed to occur in “hotspots” throughout the SN15 genome, which were often, but not exclusively, telomeric (Fig 2, Table S8). The accessory chromosome 23 (AC23) was observed to have a higher overall SNP density compared to other chromosomes. Isolates WAC2813, WAC9178, WAC2810, WAC8635, WAC13405, WAC13418, WAC13447 (all from population cluster 6) all had very few SNPs relative to SN15, forming the inner blue circle present in Fig 2. Care was taken that certain biological and technical factors did not unduly influence our interpretation of SNP density at the whole-genome level. A region on chromosome 07 of between 140,979 bp and 623,833 bp, which was identified in a previous study (Bertazzoni et al. 2021) as a potential sequencing artifact exhibited an absence of SNPs in our analysis. Low SNP counts around the rDNA tandem repeat array located on the end of Chromosome 3 (Bertazzoni et al. 2021) and other areas that overlap repeat regions may have also been caused by read-alignment depth filtering rather than the absence of variants. Repeat-induced point mutation (RIP) is an important feature indicating ‘hypermutation’ compartments throughout fungal genomes (Hane et al. 2015). The ratio of RIP-like dinucleotide changes over all transition SNPs across the pan-genome relative to the SN15 reference genome indicated that RIP-like mutations are over-represented and also tend to be localised in hotspots (Fig 3). Hotspots of RIP-like mutation tended to co-locate with regions rich in transposable elements in the SN15 genome, though there are several regions that are enriched for RIP-like mutations for a small group of isolates.

**Fig 2.**
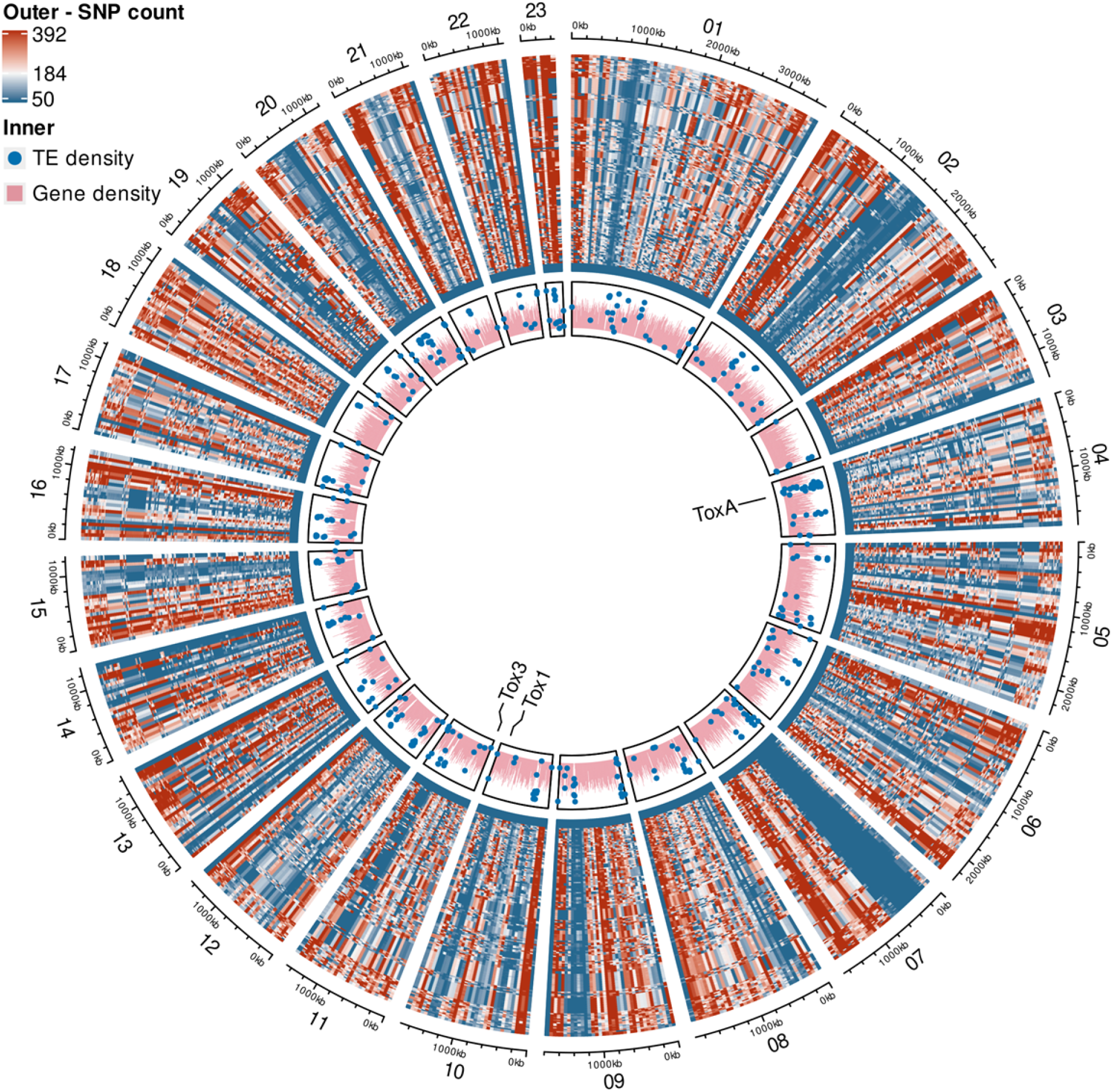
A circos plot showing SNP density over each of the 23 chromosomes in the SN15 genome assembly. The innermost track shows the proportion of bases covered by genes (CDS features, red) and transposable elements (TE, blue dots) in non-overlapping 10kb windows. For TEs, windows with TE base coverage less than 10% are not plotted. The heatmap shows SNP counts in 50 kb non-overlapping windows for each of the Western Australian isolates in the outer track (Table S8), with the colour scale boundaries set by the 10th, 50th, and 90th percentiles (50, 184, and 392, respectively).

**Fig 3.**
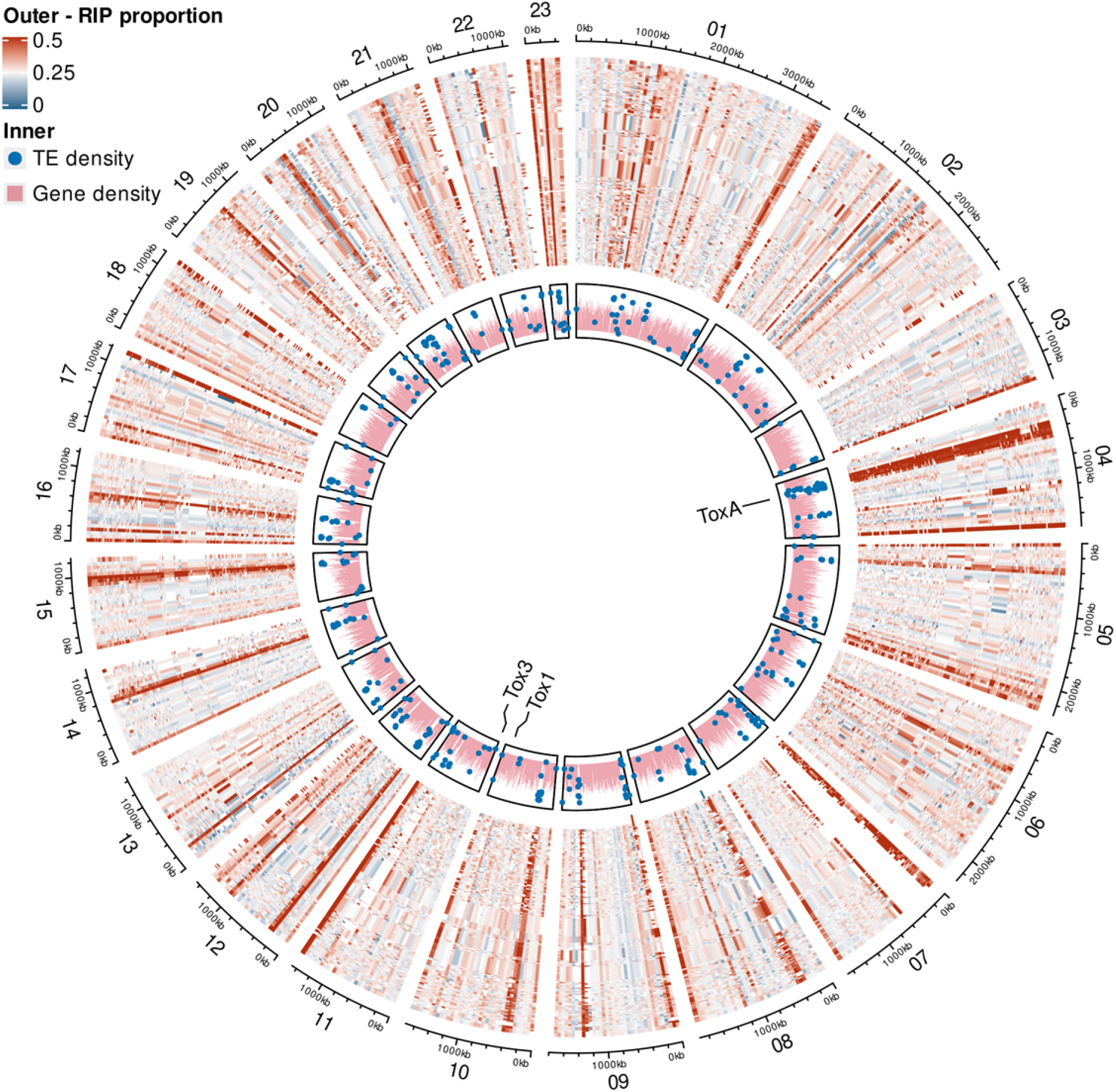
A circos plot showing the proportion of RIP-like (CA↔TA or TG↔TA) mutations over transition (C↔T or G↔A) mutations for each of the 23 chromosomes in the SN15 genome assembly. The innermost track shows the proportion of bases covered by genes (CDS features) and transposable elements (TE) in 10kb non-overlapping windows. Windows with TE base coverage less than 10% are not plotted. The heatmap in the outermost track shows the proportion of RIP-like mutations over the number of transition mutations in 50 kb non-overlapping windows for each isolate (Table S8). Windows with fewer than 20 SNPs are plotted in white to avoid high ratios caused by a small number of RIP-like mutations. By chance, 25% of transition mutations would be expected to be part of a RIP-like dinucleotide pair change.

### Comparative genomics across the WA *P. nodorum* pan-genome

The average assembly size for the 156 WA *Parastagonospora nodorum* isolates was 37.8 Mb, with a median N50 of 17 and a median L50 and NG50 of 793431 and 783556 (Table S5).

Mitochondrial assemblies produced between 1 and 3 contigs in all cases, with a median size of 49591 bp. A common ∼1000bp repeated region was observed in the mitochondrial assemblies, which appeared to be the cause of the fragmented assemblies (data not shown). The WA isolates were predicted to be highly complete with a median number of genes predicted by Genemark of 13037, with average completeness estimated via BUSCO at 98.94%, only one isolate (15FG111) had completeness estimated below 98%.

To identify potential regions of presence-absence variation (PAV), the genome assemblies of this study and previous studies (Richards et al. 2018, Phan et al. 2020) (table S1) were aligned to the SN15 reference assembly (Bertazzoni et al. 2021). In comparison to the SN15 reference assembly, the majority of the SN15 genome sequence was conserved with resequenced isolate assemblies, with small regions of PAV tending to be observed at telomere ends or large internal repeat regions (Fig 4; Table S8). In addition to being missing in isolate SN79 (Richards et al. 2018), the accessory chromosome 23 was wholly absent in isolate ‘Northam_Magenta’ and partially absent in isolates 16FG160-162, 16FG163_2, 16FG164-171, and WAC13403 (Fig 4). We observed large duplications of genomic regions in some isolates (Fig 4). On SN15 chromosome 19 there was a large duplication of a region between 850 kb and 1 Mb in length in isolates WAC13404, WAC13075, and WAC13525. The first 200 kb was duplicated in isolate Meck8 relative to SN15 chromosome 22. Isolate WAC13631 had a large duplication relative to SN15 chromosome 12, of between 750 kb and 1.1 Mb.

**Fig 4.**
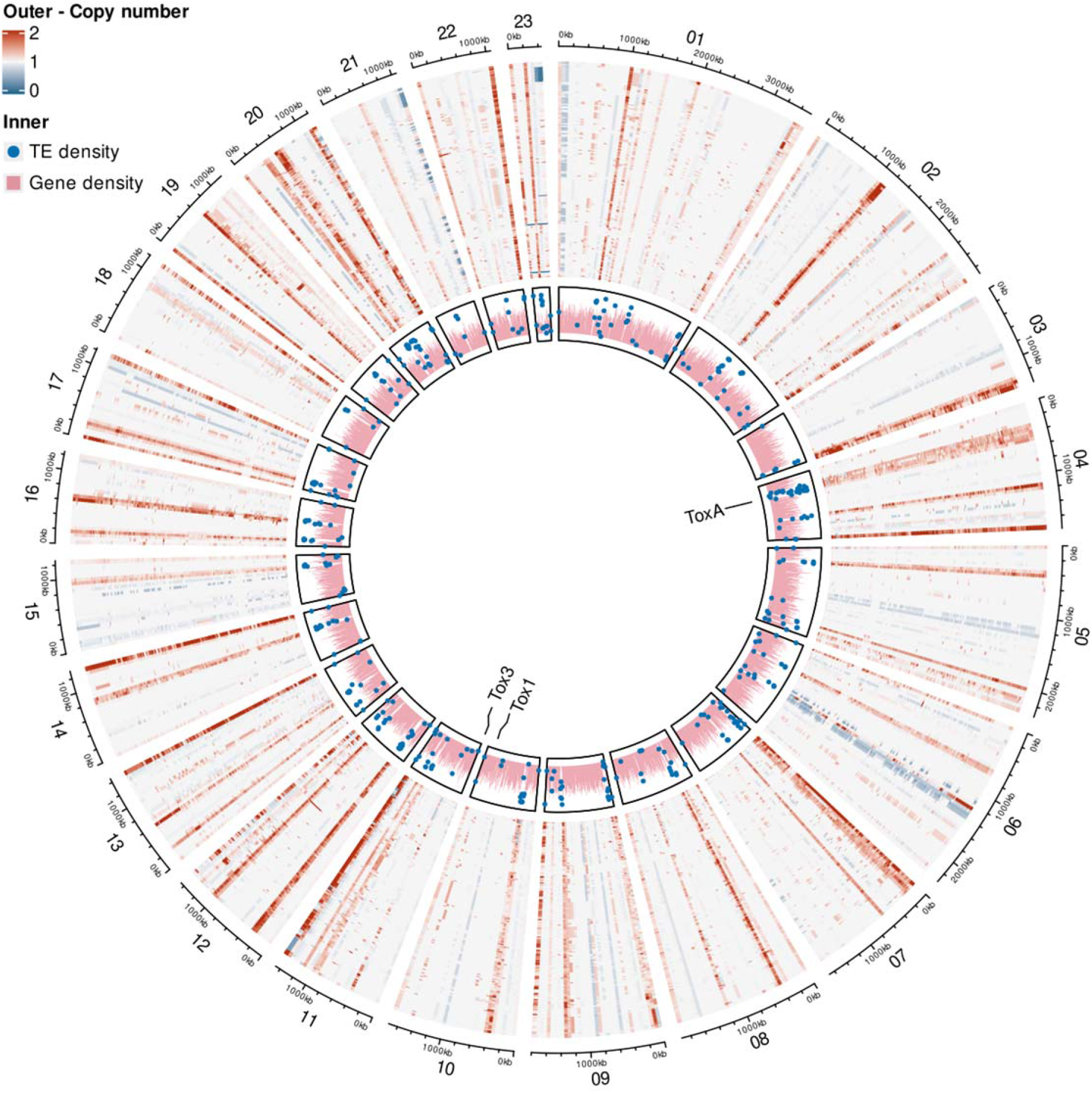
A circos plot showing each *Parastagonospora nodorum* genome assembly alignment coverage for each of the 23 chromosomes in *P. nodorum* SN15. The innermost track shows the proportion of bases covered by genes (CDS features) and transposable elements (TE) in 10kb non-overlapping windows. Windows with TE base coverage less than 10% are not plotted. The heatmap on the outside track shows average alignment coverage of each isolate genome assembly to SN15 in 50 kb non-overlapping windows (Table S8).

Genes were predicted in all genome assemblies (including previously published isolates, table S1), which resulted in a median gene count of 18294 across WA isolates, with a minimum of 17633 in isolate SN79 and a maximum of 19125 in isolate RSID03 (Table S6 and S7). The median length of coding domain sequences (CDSs) was 894 bp and was 53 bp for introns. The median BUSCO gene completeness was 3135 or 99.3%, with a median of 16 fragmented and 3 missing loci. Orthology clustering of predicted protein products from all *P. nodorum* isolates (including non-WA isolates) produced 34381 ‘orthogroups’, 14098 of which were core to the population (13628 single copy and 470 multi-copy), and 11460 were dispensable (10043 single copy, 1417 multi-copy) (Table S9). An additional 8823 singleton groups were identified (8490 single copy, 333 multi-copy) which were only observed in a single isolate. To detect orthogroups with any members potentially under positive selection at any site, dN/dS branch site tests were run for all non-singleton orthogroups using the BUSTED algorithm in HyPhy (Pond et al. 2005). This identified 5306 orthogroups that were undergoing diversifying selection at any point with p-values < 0.01. Of these, 732 orthogroups had more than 20% of sequences within the orthogroup predicted to be subject to positive selection.

Multiple codon alignment of the coding sequences of the three known effector loci *ToxA*, *Tox1*, and *Tox3* (Data S10) indicated the occurrence of non-synonymous and RIP-like mutations in these loci. The *Tox1* orthogroup (SNOO_20078A) was absent in isolates RSID36, RSID37 and RSID39 (Table S9) and was present as a single copy in all other isolates, or as a C-terminal truncated version in WAC13443 and SN79. Some branches of the orthogroup were predicted to be under positive selection (HyPhy BUSTED test, p-value >= 0.0008), and three codons showed significant position-specific positive selection (HyPhy FUBAR test, posterior probability > 0.90) at alignment codons 108 (GAC↔TCC; p=0.9194), 113 (ACC↔CCC; 0.9296), and 117 (GCA↔CAA; 0.9892). The alignment variants at T113P and R117[V,Q] are restricted to WA isolates, while the variant at D108S is restricted to a subset of the international isolates, including SN4. Overall, 13 distinct AA and nucleotide CDS sequences were detected in SNOO_20078A. The *Tox3* orthogroup (SNOO_08981AB) was present in 160 isolates, but absent in SN79, SN2000, and 8 of the 16 remaining non-Australian isolates. SNOO_08981AB was not predicted to be under positive selection (HyPhy BUSTED, p-value=0.05), and no specific positions were detected to be under positive selection using HyPhy FUBAR. Five distinct Tox3 codon sequences were observed resulting in two distinct AA sequences, with 28 WA isolates (including all isolates from population cluster 9) and one Swedish isolate (RSID28) possessing 3 mutations resulting in the codon changes N78D (AAT↔GAT), R102L (CTA↔CGA), and D104E (GAA↔GAT), with the first appearing to be RIP-like. The *ToxA* orthogroup (SNOO_16571A) was present in most isolates but absent in one WA isolate (201FG209), and 11 international isolates, including SN79. The *ToxA* orthogroup was not predicted to be under positive selection by HyPhy BUSTED, and no individual sites were predicted to be under positive selection by HyPhy FUBAR. There were non-synonymous mutations, I130V (ATT↔GTT) and E125D (GAA↔GAT), the former exclusive to WA isolates and the latter being RIP-like and present in seven isolates, of which 2 were from WA. Five anomalous ToxA CDS sequences were observed, each with distinct AA sequences, however all of the predicted genes lacked a C-terminal region (which included the RGD motif) annotated in SN15, possibly indicating exon mis-annotation in these isolates.

Effector prediction using Predector (https://github.com/ccdmb/predector) identified 779 effector candidates in the SN15 reference isolate which were predicted to be secreted and with positive EffectorP 2 scores, and 1348 effector candidates with Predector scores greater than zero, of which 132 were homologous to known fungal effectors or had virulence associated Pfam domains. Across the entire pangenome, 2055 orthogroups were predicted to be secreted and have a positive EffectorP 2 prediction, 3398 orthogroups had members with a Predector effector score greater than zero. Of the predector candidates 997 were predicted to be secreted and +have a positive EffectorP 2 prediction, 145 contained effector homologues or virulence related Pfam domains, 411 had members significantly under positive selection, 55 were under positive selection in more than 20% of orthogroup sequences, 750 were accessory orthogroups, and 1405 were singleton orthogroups.

Orthogroups showed some PAV spanning large regions of the genome (Fig 5, Table S9). The largest of these was only present in 31 isolates and contained 385 orthogroups, comparable in gene number to AC23 which contains 218 predicted genes in SN15. Assembled scaffolds containing orthogroups in this PAV group were generally shorter than 200 kb, did not align to any chromosomes in the SN15 genome, but did align to scaffolds in other isolates with a high level of collinearity (Data S11).

**Fig 5.**
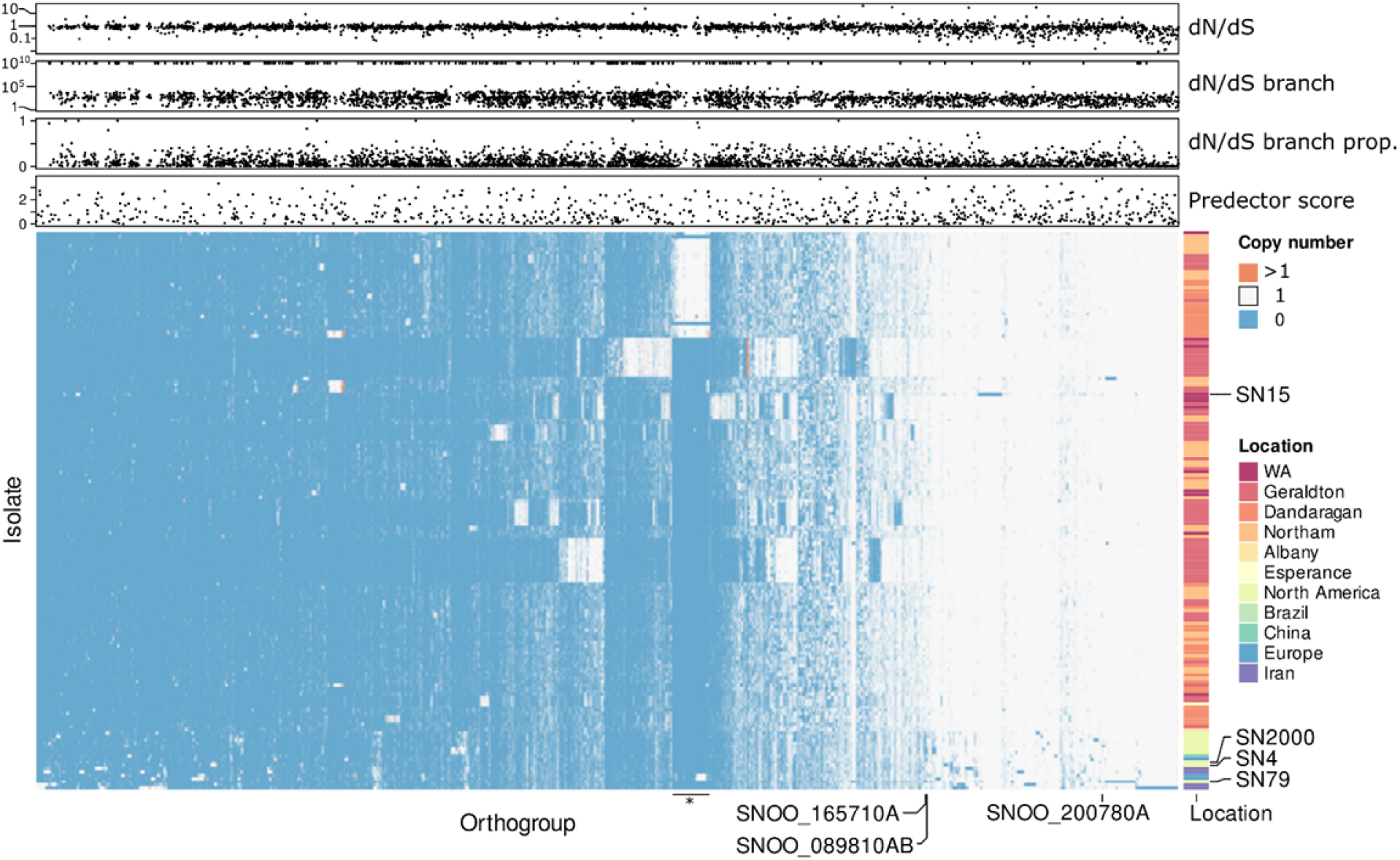
Dispensable and multi-copy orthogroups for each isolate in the *P. nodorum* pan-genome. “Orthogroups” is used as a general term to cover both orthologues and paralogues, which have not been separated here. Heatmap rows represent each *P. nodorum* isolate, and columns indicate each of the dispensable or multicopy orthogroups. Heatmap colour indicates the number of copies of an orthogroup each isolate has. Orthogroups absent (blue), present with single copy genes (white) and present with multicopy genes (orange) are shown. The columns of orthogroups containing ToxA (SNOO_16571A), Tox3 (SNOO_08981AB), and Tox1 (SNOO_20078A) are indicated. Locations that isolates were collected from are indicated on the right hand side colour bar. The rows corresponding to reference *P. nodorum* isolates are indicated. The top 3 scatter plots indicate orthogroups with any members with significant positive selection tests (p < 0.01). “dN/dS” indicates the overall dN/dS for the whole orthogroup. “dN/dS branch” indicates the dN/dS at the branch predicted to be under the highest selection. “dN/dS branch prop” indicates the proportion of sequences in the orthogroup predicted to be under positive selection. “Predector score” indicates where the highest scoring member of an orthogroup was greater than 0.

### Accessory regions and candidate effector loci are enriched in RIP-like mutations and unknown functions across the WA pan- genome

Statistically significant enrichment (two-tailed hypergeometric test, BH FDR corrected p-value < 0.05) of predicted protein functions (by gene ontology (GO) terms) were observed within various subsets of pan-genome orthogroups. The core pan-genome was enriched for 3509 GO terms generally associated with core functions including biosynthesis, cell cycle control, and transport (Table S11). Conversely, the accessory pan-genome was depleted in 1698 terms associated with core functions. Similarly, singleton orthogroups (present in only one isolate) were depleted in 1708 GO terms associated with core functions, as were orthogroups not present in the SN15 reference isolate (2946 depleted). Multicopy subsets of the core, accessory, and singleton pan- genomes, and the subset of the accessory pan-genome containing between 20% and 80% of isolates showed similar patterns of enrichment and depletion as their respective supersets. Orthogroups where more than 20% of sequences were predicted to be positively selected were depleted in 139 GO terms relating to core functions (e.g. transport, metabolic processes, response to stimulus).

Genes of the SN15 reference isolate that were RIP-affected (above the RIP ratio 95th percentile, 0.432) were depleted in 54 core GO terms. AC23 contained two of the four genes in SN15 predicted to be involved in dehydroaustinol biosynthesis, but otherwise GO terms were all depleted. The large PAV group was depleted in 32 terms with general core functions. Overall, the enrichment tests above did not reveal any clear associations between GO terms and features within the genomic landscape. The majority of orthogroups (28767 of 34381) had no GO terms assigned. A complementary series of enrichment tests for a lack of GO terms (‘no-GO’) were performed using fishers exact tests with an uncorrected p-value threshold of 0.05. The core pan-genome was depleted for no-GO orthogroups, whereas the accessory and singleton pan-genome, positively selected orthogroups with more than 20% of sequences under selection, the PAV group, RIP-affected orthogroups and SN15 loci on AC23 were all significantly enriched no-GO orthogroups.

Enrichments tests were also performed for predicted secreted proteins and effector-like proteins (predicted secretion and EffectorP2). The accessory pan-genome, singleton pan-genome, orthogroups absent in SN15, and positively selected orthogroups were enriched for secreted proteins. However the core pan-genome and PAV groups were depleted in secreted proteins. Similarly the accessory pan-genome and positively-selected orthogroups were enriched in effector-like orthogroups, while the core and singleton pan-genomes were depleted.

To find effector candidates a representative member of each orthogroup for each distinct locus was selected (Table S10). For orthogroups with members in the reference isolate SN15, all distinct loci were included selecting the protein isoform with the closest sequence length to the average orthogroup length. Representative members of other orthogroups were selected by taking the member with the closest sequence length to the average orthogroup length, with a preference for alternate reference isolates SN4, SN2000 and SN79. Orthogroups with a Predector score greater than zero, or with a signal peptide predicted by any method and an EffectorP 2 score greater than 0.5 were selected to be effector candidates. This identified 3579 candidate orthogroups, of which 788 and 1504 were in the accessory and singleton pangenomes, respectively. The WA isolates contained 1362 candidates (181 accessory) that were not predicted in the international isolates, of which 411 were restricted to the non-core populations (not cluster 4; 64 accessory). The core WA population (cluster 4) possessed 842 distinct candidates not present in other clusters (64 accessory), while clusters 6 (which includes SN15) and 8 (international) possessed 96 (1 accessory) and 375 (52 accessory) unique candidates. The reference isolates SN15, SN4, SN2000, and SN79 were missing 1732 candidate orthogroups (317 accessory).. There were 66 candidate orthogroups predicted to be under positive selection in at least 20% of orthogroup members, including a Tox3 homologue (SNOO_01097A). The large clusters of orthogroups with PAV identified in Fig 5 contained 18 candidate orthogroups. From these 3579 candidates, 1809 orthogroups with known functions or similarity to known effectors were selected (Table 2. Table S10).

**Table 2.**
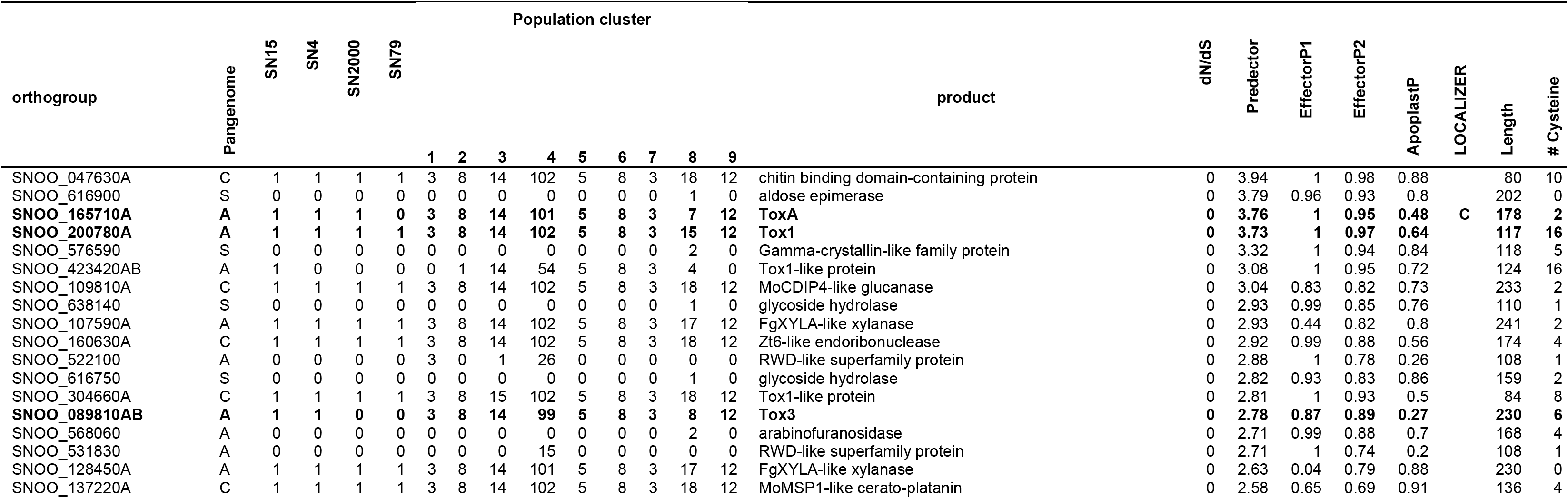
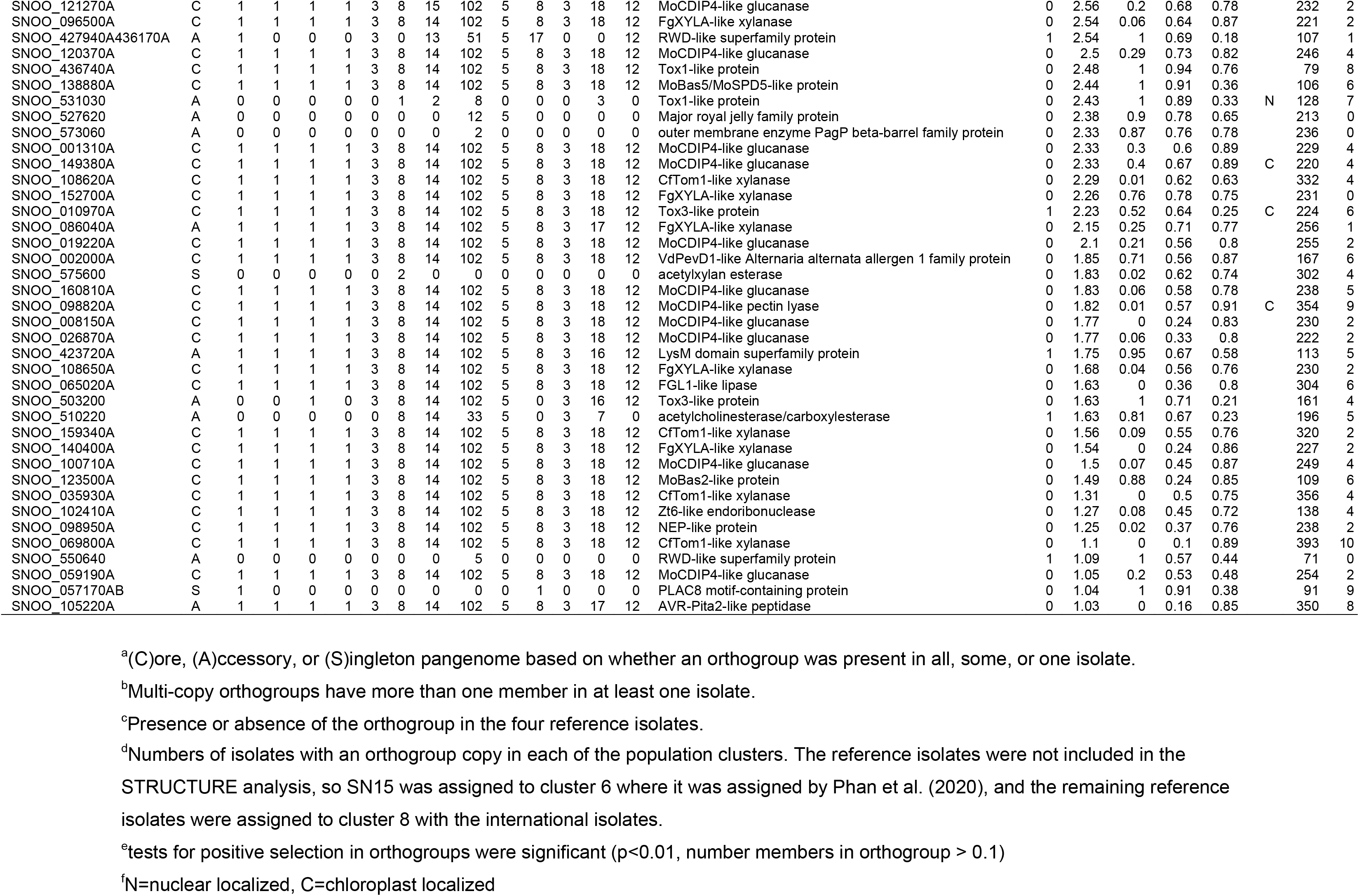
Selected effector candidate orthogroups with functional annotations in the *P. nodorum* pangenome. Of the 1908 effector candidates with functional predictions, this table includes all candidates with effector homologues and a Predector score > 1, the 5 top ranking (by Predector) orthogroups that were under positive selection, the top 5 candidates specifc to any populations identifed by structure, and the top 5 candidates restricted to Western Australian or non-core (satellite) WA population clusters. Population cluster 8 includes the international isolates, and cluster 4 represents the core WA cluster. For each candidate orthogroup SN15 members were selected as representatives, or in the case of orthogroups absent in SN15 the member with the highest Predector score. Membership of PAV clusters shown in Fig 5 is indicated by the corresponding number, where applicable. Products in bold show where a candidate matched a known necrotrophic or avirulence effector. All orthogroup candidates presented here have a signal peptide predicted by at least one method, and none any predicted transmembrane (TM) domains.

Tox1 (SNOO_200780A, SNOO_304660A, SNOO_423420AB, SNOO_436740A, SNOO_531030), four Tox3 (SNOO_089810AB, SNOO_010970A, SNOO_438650A, SNOO_503200), and one ToxA (SNOO_165710A) homologous orthogroups were identified, including each of the original sequences from the SN15 isolate. Numerous other effector homologues were found, including 19 MoCDIP4 (Chen et al. 2013), 7 XYLA (Pollet et al. 2009, Sperschneider et al. 2015), CfTom1 (Pareja-Jaime et al. 2008, Ökmen et al. 2013), 2 FGL1 (Voigt et al. 2005), 2 AVR-Pita/AVR-Pita2 (Dai et al. 2010, Chuma et al. 2011), 2 Zt6 (Kettles et al. 2018), 2 HCE2/Ecp2 (Stergiopoulos et al. 2012), 2 NEP/NLP (Oome et al. 2014), 2 BEC2 (Schmidt et al. 2014) and 5 other CFEM domain containing proteins, 2 Cgfl (Fungalysin peptidase) (Sanz-Martín et al. 2016), and one each of BEC1019 (Zhang et al. 2019), NIS1 (Yoshino et al. 2012, Irieda et al. 2019), MoBas2 (Mosquera et al. 2009), MoCDIP1 (Chen et al. 2013), MoMSP1 (Wang et al. 2016), MoSPD5/MoBas4 (Mosquera et al. 2009, Sharpee et al. 2017), PevD1 (Bu et al. 2014), and ZtNIP2 (M’Barek et al. 2015). Other notable functions and families identified among these effector candidates include peptidases, nucleases, cupredoxins, CAP-superfamily proteins, Egh16-like virulence factors, Osmotin/Thaumatin-like proteins, Killer toxin KP4, tuberculosis necrotizing toxin, SnoaL-like/NTF2-like domain superfamily, RmlC-like cupin domain superfamily, TolB-like/major royal jelly protein, Ubiquitin and biotin related functions, and WD40/Ankyrin/Kelch repeat-containing proteins. A single UstYa-like mycotoxin biosynthesis protein, and several proteins related to metabolite biosynthesis or detoxification were also found.

## Discussion

The transition from the sequencing of a single or few reference genomes to larger populations has broadened the scope of comparative genomics of plant pathogens. This includes the identification of: additional accessory genome content missing from the reference isolate (Badet and Croll 2020); spatial distribution of virulence loci, and; region-specific selection pressures (Richards et al. 2019). To this end, we investigated the interplay between the population structure and genomic features relevant to plant pathogenicity, in a Western Australian population of *Parastagonospora nodorum*.

### Population structure shows distinct regional sub-populations

Previously we have analysed the Western Australian *P. nodorum* population using SSR markers (Phan et al. 2020), and identified five distinct clusters of isolates. Two main clusters were proposed to represent a gradual change over time in response to wheat cultivar use, while the three remaining clusters were highly similar and were proposed to be clonally expanded populations. Interestingly, although the two core populations had a 1:1 ratio of mating type loci, the core population was observed to be in linkage disequilibrium, suggesting a predominantly asexually reproducing population. In contrast, this study indicated that the WA *P. nodorum* population is dominated by a single diverse main population, with seven satellite clusters that are highly similar. The fact that the core population was not split into two as previously observed may be explained by the marker type and number, the number of clusters selected to find, and the clustering method employed, where STRUCTURE (used in this study) explicitly models gene flow. The relative uncertainty of individual assignment to cluster 4 (the main population), suggests that there is some latent structure in the data, but this appears not to correspond to distinct reproductively isolated populations. In addition to the three satellite clusters identified by Phan *et al*. (Phan et al. 2020), the greater resolution gained by the use of many SNPs identified an additional 4 minor clusters, which were subdivided from the two previously identified core clusters (Fig S3). The relatively low diversity observed in these seven clusters suggests that these are very recent expansions or clonal subpopulations. Analysis of the index of association indicated that all but two small sub-populations are in linkage disequilibrium, suggesting that limited sexual reproduction is occurring in populations. It should be noted that the two exceptional population clusters (1 and 7) contained few isolates and their lack of linkage disequilibrium may be inaccurate. Linkage disequilibrium across WA *P. nodorum* sub-populations was also reported by Phan *et al*. (Phan et al. 2020), but they observed that clusters P1 and P2 (corresponding to cluster 4 in this study) had a 1:1 mating type ratio which indicates the potential for sexual reproduction, and that cluster P5 (corresponding to cluster 9 in this study) was in linkage equilibrium. The finding that the international cluster is in linkage disequilibrium is also unexpected as it would imply that the global *P. nodorum* population is largely asexual, in conflict with numerous previous reports (Keller et al. 1997, Caten and Newton 2000, Murphy et al. 2000, Sommerhalder et al. 2006, Stukenbrock et al. 2006).

Although the r_d_ values are significantly different from a random background, they are still relatively low so the population may be exhibiting a mixture of clonal and non-clonal reproduction. Alternatively, the permutation method of r_d_ may not be appropriate for samples with large numbers of SNPs in relatively high density (compared to low-throughput marker studies). Phan *et al*. (Phan et al. 2020) suggested that the non-core populations may be hybrids of local and internationally introduced *P. nodorum* isolates. This is not supported by the phylogenetic tree or population structure analysis reported in this study; however, STRUCTURE analysis indicated a cluster of six isolates from population 4 collected from the Northam and Dandaragan regions which had a high posterior probability of assignment to the international subpopulation, suggesting possible historic exchange of genetic material.

In the overall WA population there was no correlation observed between sampling location and genetic distance when nearly clonal isolates were excluded, suggesting that there are no geographic barriers to migration. Similarly, PCA only indicated an axis of variance separating international isolates from those from WA, with no other principal components showing association with sampling location or year (Fig S6 and S7). Long range wind dispersal of sexual ascospores has long been known to occur in the Western Australian *P. nodorum* population (Bathgate and Loughman 2001), though dispersal by infected seed is often reported internationally (Cunfer 1978, Cunfer 1998, Bennett et al. 2005). Both of these mechanisms may explain the lack of geography-dependent population structure observed in this study. Although seed borne dispersal is more likely if the population is mostly asexual, we were unable to find publications describing seed borne epidemics of *P. nodorum* in Australia.

The majority of isolates from the satellite clusters were collected from northern regions of the WA sampling area, predominantly Geraldton and Mingenew. Within the sampling zones presented in this study, the average rainfall in the south west regions is typically higher than in the dryer northern regions (http://www.bom.gov.au/climate/current/annual/wa/summary.shtml).

In addition to splash dispersal of secondary inoculum and a general positive correlation of rainfall with *P. nodorum* disease load (Solomon et al. 2006, Shaw et al. 2008), rain impacts can indirectly enable long-distance air travel (Kim et al. 2019), and may have contributed to the increased diversity of south-western regions. Similarly, high temperature during harvest time has been observed to be negatively correlated with *P. nodorum* disease load (Shaw et al. 2008), which may favour stronger populations in the southern regions. Numerous environmental factors can influence the lifecycle of *P. nodorum* which may explain these northern clonal sub- populations. However, the samples used in this study were not collected with population genetics analyses in mind, and an intentionally designed experiment may yet reveal the existence of similar structure in the other regions of the WA population. It appears that there is variance in the WA population, but this remains cryptic and is not explained by barriers to gene flow.

The presence of necrotrophic effector loci, or in some cases specific allele variants, is a direct determinant of crop disease outcomes in combination with the corresponding host sensitivity loci (Vleeshouwers and Oliver 2014). A US-based pan-genome study previously indicated alternate sets of candidate effector loci between two major *P. nodorum* sub-populations (Richards et al. 2019), highlighting the importance of region-specific genomic analysis and refinement of effector predictions in local isolates. The widespread surveillance of effector profiles within pathogen populations has great potential for crop disease management tailored to specific regions. In this WA-based study, low diversity sub-populations tended to have a conserved haplotype profile for the 3 known effector loci *ToxA*, *Tox1* and *Tox3* (Fig 1). In contrast sub-population cluster 5 exhibited notable diversity in its Tox1 effector haplotypes. Overall this study indicates the potential for resistant cultivars to be broadly recommended for growing across regions identified to have low diversity, but less reliably for regions with higher diversity. Given the extreme potential for genome plasticity in fungal genomes (McClintock 1941, Hane et al. 2011, Croll et al. 2013, Testa et al. 2016), it is encouraging that conserved effector haplotype profiles were observed in several cases. Identification of geographic regions exhibiting high variability of effector haplotype profiles within a narrow timeframe may also become an important element of crop disease monitoring in the future.

### Pan-genome comparisons highlight regional diversity across the genomic landscape and effector contents

A previous comparison of *P. nodorum* isolates from the USA and Australia (Richards et al. 2018, Bertazzoni et al. 2021), indicated that the smallest chromosome (AC23) is an accessory chromosome with high levels of mutation, diversifying selection, and numerous gene duplications for redundant pathogenicity-related functions. Both AC23 and the region of chromosome 4 encoding the known effector gene *ToxA*, appeared to exhibit structural mutations that may be influenced by breakage-fusion bridge (BFB) formation. BFBs are hypothesised to be a driver of accessory chromosome formation and evolution (Croll et al. 2013, Bertazzoni et al. 2018) and of intrachromosomal recombination events that cumulatively lead to an inter-species conservation pattern termed “mesosynteny” (Hane et al. 2011). The largest duplications or absences in WA isolates relative to SN15 were predominantly located near telomeric regions or on AC23 (Fig 4). This is consistent with the known enrichment of structural rearrangement in subtelomeres (Hocher and Taddei 2020) and proposed prevalence of BFB-mediated rearrangement across the Dothideomycetes (Croll et al. 2013). Complete and partial absences of SN15 AC23 were observed in some Western Australian isolates, suggesting that large structural mutations are occurring in the field (Fig 4).

SNP density across the WA pan-genome was also consistently highest on AC23 (Fig 2) and near telomeres; however, several intrachromosomal mutation hotspots were also observed, where overall gene and repeat densities appeared relatively normal. The three known effector loci *ToxA*, *Tox1* and *Tox3* are all located at or near telomeres in SN15 (Bertazzoni et al. 2021), which also corresponded with SNP hotspots. Additionally a large orthogroup PAV cluster was observed in a subset of WA isolates, which may represent a previously undescribed accessory chromosome. Future long-read sequencing of these isolates may resolve the structure and history of these fragmented regions.

Overall, the variable genes and genome regions across the pan-genome (PAV and SNP) did not highlight any clear association with known gene functions. Indeed, the gene ontologies resource used in this study define function very broadly, and are biased towards conserved functions. Variable regions, including accessory, diversifying, repeat-rich and RIP-mutated, appeared to be bereft of known gene functions, but were conversely enriched in effector-like candidate loci. We observed 181 effector candidates that were absent in the international isolates and present in more than one WA isolate, of which 68 were restricted to a single WA subpopulation. This suggests that WA may have a distinct pathogenicity gene profile; however, the number of non-Australian isolates used in study was relatively small, so further comparison with international isolates may find these candidates elsewhere. The large PAV cluster of orthogroups which may represent an accessory chromosome or large chromosomal PAV, contained 18 effector candidates, including two putative cupredoxins. Five Tox1 and four Tox3 homologous orthogroups in the pan-genome were found in different frequencies across the different populations. This suggests that these NEs may share a common origin but have since duplicated and diversified sufficiently that they were predicted as distinct orthogroups. Expansion and diversification of effectors within pathogen genomes appears to be a common phenomenon (de Guillen et al. 2015, Praz et al. 2017) in plant pathogens, and these homologues are strong effector candidates which may confer distinct phenotypes from their characterised homologues. Numerous homologues of effectors from other species were also identified, of which the necrotrophic effectors Zt6 and ZtNIP2 homologues are of particular interest to *P. nodorum*. Zt6 is a ribotoxin which cleaves non-self ribosomal sarcin-ricin loops in both wheat and microbial competitors (Kettles et al. 2018). ZtNIP1 induces light-dependent necrosis in wheat with differential responses between cultivars (M’Barek et al. 2015). Interestingly, numerous other homologues of avirulence elicitors and biotrophic effectors were also identified. Many of these have functions related to nutrient and sugar scavenging, but may still have a plausible virulence role in the necrotrophic *P. nodorum*. For example, numerous MoCDIP4 homologues were identified which induces cell death in non-host plants of *Magnaporthe oryzae* (Chen et al. 2013) and interferes with mitochondrial homeostasis, which in turn inhibits mitochondrially mediated resistance responses (Xu et al. 2020). The MoCDIP4-like effector candidate SNOG_01146 (SNOO_01146A) is significantly upregulated *in planta* and in a mutant lacking the effector-regulating PnPf2 transcription factor (Jones et al. 2019). Other effector-like orthogroups without fungal effector homologues but with potentially virulence-related functions were also identified. The putative aldose epimerase SNOO_61690 was one of the highest ranked candidates and only observed in isolate RSID03, but appears to be a highly truncated copy of SNOO_063350A which is present in all other isolates. In *Phytophthora sojae* the apoplastic aldose 1-epimerase AEP1 functions as a virulence factor by scavenging apoplastic aldose, and triggers cell death and pattern triggered immunity in *Nicotiana benthamiana* (Xu et al. 2021).

Comparative genomics across the WA pan-genome also indicated the prevalence of RIP-like mutations in variable regions, and was also associated with candidate effector loci. The presence of distinct genome wide patterns of RIP-like mutation between isolates would indicate that RIP has been actively occurring within a recent time frame. However RIP-like mutations did not correlate with diversifying selection observed across the whole genome, or even within highly ranked effector loci. RIP occurs during pre-meiosis, however, the observation of linkage disequilibrium suggests that the WA population may not be regularly undergoing sexual recombination. Although previous studies indicate *P. nodorum* has the potential for meiosis in WA (Murphy et al. 2000, Bathgate and Loughman 2001), it is yet to be determined if widespread RIP has only occurred in the past or if recent selection pressures have eliminated background isolate diversity. Investigations of the role of RIP in biotrophic and hemibiotrophic pathogens (Testa et al. 2016, Gervais et al. 2017) have indicated that RIP-mediated loss of a recognized avirulence effector may confer a selective advantage. We speculate that the nature of necrotrophic effectors with inverse gene-for-gene interactions with host sensitivity receptors (Fenton et al. 2009, Thrall et al. 2016) may conceal the full influence of RIP in necrotrophic effector diversification, as loss of function mutations are unlikely to be advantageous and would be selected against in the population. The remaining detectable RIP would be observable only for genome regions which do not significantly contribute a selective advantage. Nevertheless the *ToxA* locus resides in a large RIP hotspot on chromosome 4, and the confirmed effector loci *ToxA*, *Tox1* and *Tox3* retain a small number of non-synonymous RIP-like SNPs. The infrequent but constant potential for RIP to introduce potentially virulence enhancing mutations remains an important consideration for genome-guided disease risk assessment in necrotrophs.

## Conclusion

Population-level pan-genome approaches are the next frontier of plant pathogen bioinformatics, which may eventually lead to affordable genome-based crop disease diagnostics and surveillance at a local level. Trends in pathogen genomics have begun to abandon intensive study of a single reference isolate, and are steadily progressing towards regionally-customised and data-driven assessments of pathogen gene-content, particularly with regards to effector genes (Richards et al. 2019, Badet and Croll 2020, Bertazzoni et al. 2021). In this study, we analyse a local Western Australian population of the wheat pathogen *Parastagonospora nodorum* and identify multiple genome features of relevance to this pathosystem. We observed an apparently high potential for genome adaptability, suggested by the presence of active RIP and other mutations, but this was not readily observed to drive diversification of its three known highly conserved necrotrophic effectors. Mutation hotspots were identified which were rich in effector candidates and genes of unknown function, and often also classified as dispensable, sub-telomeric or large repeat-rich regions. In a spatial context, we observed regional ‘hot’ and ‘cold-spots’ of population diversity, that may be linked to climatic factors affecting spore dispersal. Across the local pan-genome, we observed the diversity of haplotype profiles of 3 known effector genes to be conserved in regions with lower overall diversity. A total of 3579 novel effector candidates were predicted across all isolates, with 2291 of these exhibiting PAV across the genomes and 1362 restricted to WA isolates. Overall this study has progressively improved bioinformatic resources for the *P. nodorum* pathogen, as well as advancing approaches for the study of fungal pan-genomes with a view towards developing a region-specific understanding of host-pathogen interactions.

## Methods

### DNA extraction and sequencing of Western Australian *P. nodorum* isolates

Genomic DNA of 141 previously described Western Australian *P. nodorum* isolates (Phan et al. 2020) were extracted (Xin and Chen 2012) and sequenced by the Australian Genome Research Facility (Melbourne, Australia) (Illumina HiSeq2500, TruSeq PCR-free, 125bp paired end (PE), 600 bp insert size) [NCBI BioProject: PRJNA612761] (table S1). Genomic DNA of 17 new isolates and 14FG141 and Mur_S3 of the previous 141 were extracted using a Qiagen DNeasy Plant Mini kit (Venlo, Netherlands. Catalogue ID: 69104) and sequenced by Novogene (Beijing, China) (Illumina HiSeq2500, TruSeq PCR-free, 150bp PE, 350bp insert size). Pre-existing draft genomes of 15 international *P. nodorum* isolates (Syme et al. 2018) [NCBI BioProject: PRJNA476481] and chromosome-scale genome assemblies for the reference isolate SN15 (Bertazzoni et al. 2021), and for US isolates SN4, SN2000 and SN79-1087 [NCBI BioProject: PRJNA398070] (Richards et al. 2018) were also used.

Reads were trimmed for TruSeq universal adapters and low-quality using CutAdapt (v1.18) (2 passes, 3 trims/pass, terminal Phred score >2, average Phred score ≥5, length ≥50) (Martin 2011), and for contaminants (e.g. PhiX) using BBduk (v38.38, read kmer coverage of 0.7) (https://jgi.doe.gov/data-and-tools/bbtools/bb-tools-user-guide/bbduk-guide/), using the UniVec database (https://www.ncbi.nlm.nih.gov/tools/vecscreen/univec/) and the PhiX genome (NCBI RefSeq: <u>NC_001422.1</u>) (Sanger et al. 1978) as bait templates. Potential sample contaminants were searched for using Kraken (version 2.0.7) (Wood et al. 2019) searching against a database constructed from all NCBI Refseq bacterial, archaeal, protozoan, viral, and fungal genomes (downloaded: 2019-03-16), as well as the human GRCh38 genome (Wood and Salzberg 2014) (table S2). The four published reference *P. nodorum* genomes (Richards et al. 2018, Bertazzoni et al. 2021) were also included as a positive set. Reads were aligned to the four reference *P. nodorum* genomes using BBmap (https://jgi.doe.gov/data-and-tools/bbtools/bb-tools-user-guide/bbmap-guide/, version 38.38) to evaluate insert size and completeness. Quality control statistics for each step were collected using FastQC version 0.11.8 (https://www.bioinformatics.babraham.ac.uk/projects/fastqc/), BBmap and Samtools (Li et al. 2009), and were collated using MultiQC (Ewels et al. 2016) (Data S1). Code for performing QC steps is available on GitHub at https://github.com/darcyabjones/qcflow (commit: <u>e77715d</u>).

### Read alignment and variant calling

Short reads from all samples were aligned to the *P. nodorum* SN15 reference genome (Bertazzoni et al. 2021) using Stampy version 1.0.32 allowing multi-mapping reads (-- sensitive --substitutionrate=0.0001 --xa-max=3 --xa-max-discordant=10) (Lunter and Goodson 2011). PCR duplicate reads in the aligned binary alignment map (BAM) files were marked using Picard version 2.18.29 (http://broadinstitute.github.io/picard/). Short variants were then predicted from the BAM files using GATK version 4.1.0.0 (McKenna et al. 2010, Poplin et al. 2018) and the following “bootstrapped” pipeline: 1) Call individual variants using gatk HaplotypeCaller. 2) Combine variants from all samples and run joint genotype prediction using gatk GenotypeGVCFs. 3) filter first pass variants with extreme statistics for each mutation type (snp, indel, or mixed) using gatk VariantFiltration. 4) Recalibrate the base quality scores in the BAM files using the predicted variants, using gatk BaseRecalibrator and gatk ApplyBQSR. 5) Steps 1-4 were repeated using the recalibrated BAM files until there was no difference in base quality score recalibration (BQSR) statistics between successive iterations.

Filters applied at each bootstrap iteration are detailed in Table S3. An initial set of variants was found from the 140 isolates initially sequenced (excluding the resequenced SN15 isolate), using 5 bootstrap iterations to converge the BQSR scores. Nineteen additional sequenced isolates and 18 previously sequenced isolates (Syme et al. 2018), were then included in the analysis using the previously identified variant loci as starting points for two further bootstrap iterations including alignments from all isolates. Unfiltered variants were taken from the final bootstrap step and filtered more stringently. Individual genotype depth and genotype quality scores were visualised using the R (Team 2013) packages vcfR (Knaus and Grünwald 2017) and ggplot2 (Wickham 2016) to determine appropriate statistic cutoff thresholds, and were soft filtered using gatk VariantFiltration to have a minimum genotype “DP” statistic of 8 and minimum “GQ” score of 30. Variant locus “DP” and “MQ” score ranges to include were determined based on scores in non-repetitive regions visualised using vcfR chromoplot. Other variant loci statistics were visualised using ggplot2 to determine cutoff thresholds, and loci were filtered using gatk VariantFiltration separately for each variant type using the selected thresholds. All filtering parameters used during variant prediction are presented in table S3. Filtered variants with effects on the translation of annotated genes (Bertazzoni et al. 2021) (i.e. non-synonymous or nonsense mutations) were identified using SnpEff (Cingolani et al. 2012).

### Phylogeny estimation and population structure analysis

Single nucleotide polymorphisms (SNPs) for phylogenetic and population analyses were selected by excluding SNPs missing in more than 30% of isolates or with a SnpEff impact prediction of “HIGH” or “MODERATE”. To reduce the potential impact of small scale linkage disequilibrium, SNPs were selected using PLINK version 1.9 (Purcell et al. 2007) by taking SNPs with the highest minor allele frequency where two SNPs within 10 kb have an R^2^ value greater than 0.6 (--indep-pairwise 10 kb 1 0.60). Resulting SNPs were then filtered to have a minimum non-major allele frequency of 0.05 using BCFtools (Li 2011) (--min-af “0.05:nonmajor”). SNPs were converted to a sequence alignment and the substitution model best fitting the data was predicted using ModelFinder in IQTree version 2.0.3 (-st DNA -m “MF+ASC” -mset “GTR,JC,F81,K80,HKY,K81” -cmax 15 -rcluster 25 -safe) (Kalyaanamoorthy et al. 2017). The best performing model was selected and used to construct a maximum likelihood tree using IQTree with 10000 UFBoot (Hoang et al. 2018) and 1000 SH-aLRT iterations (-bb 10000 -bnni -alrt 1000 -st DNA) (Minh et al. 2020). Phylogenetic trees were plotted using the R version 4.0.2 packages phytools v0.7-47 and ggtree v2.2.4 (Yu et al. 2017). Phylogenetic trees from this study and the tree published by Phan *et al*. (Phan et al. 2020) were visually compared using tanglegrams, implemented in the dendextend (version 1.14.0) package (Galili 2015).

Population structure was inferred from SNP data using STRUCTURE version 2.3.4 (Pritchard et al. 2000). To select an appropriate number of subpopulations (K) to model, STRUCTURE was run with 10000 replicate burn-in period and 20000 MCMC replicates for a range of values of K between 1 and 12, rerunning 8 times for each K to account for random starting points. The optimal value of K was selected using STRUCTURE HARVESTER (Earl and vonHoldt 2012) using the method described by Evanno *et al*. (Evanno et al. 2005). STRUCTURE was run using the selected value of K, using a 20000 replicate burn-in period and 100000 MCMC replications, running with 8 random seeds and selecting the run with the highest log probability of data given the model.

Population statistics were calculated and visualised using R version 4.0.2 packages: ade4 v1.7- 15 (Thioulouse et al. 1997), adegenet v2.1.3 (Jombart 2008), poppr v2.8.6 (Kamvar et al. 2014), and vcfR v1.12.0 (Knaus and Grünwald 2017). The filtered SNP variants were imported into R and locus heterozygosity and G′_st_ (Hedrick 2005) scores were computed using vcfR. To account for effects near identical multilocus genotypes in population statistics calculations, the maximum likelihood (ML) distances from IQTree were used to identify MLGs that are highly similar and collapse them using poppr. For each cluster identified by STRUCTURE, the Shannon, Simpson, and inverse Simpson indices (Hurlbert 1971) were calculated, for both MLG collapsed and uncollapsed data using poppr. To test for linkage disequilibrium in population clusters, isolates from each population cluster were selected and a permutation test of *r_d_* (a modification of *I_A_*) (Agapow and Burt 2001) with 999 permutations was run for polymorphic loci within those isolates using poppr. To test for any correlation of geographic distance with phylogenetic distance in the WA isolate population, a Mantel test with 999 replications was performed using ade4 comparing ML distances from IQTree with euclidean distances of GPS coordinates from Australian clone corrected isolates with known sampling locations and distinct collapsed multilocus genotypes. Patterns of variance were assessed using principal component analysis (PCA) of the filtered SNPs from all isolates with distinct collapsed MLGs using the adegenet package.

### Genome assembly

Overlapping 150 bp paired end reads from the WA isolates were stitched using BBmerge version 38.38 (Bushnell et al. 2017) using the strict mode, kmer size of 62 bp, allowing assembly extension up to 50 bp from the ends of reads (rem mode), and using error correction to assist merging (ecct option). Genomes were assembled using Spades version 3.13.0 (Bankevich et al. 2012) (parameters: --careful --cov-cutoff auto). Different kmers were used depending on input read length (Supp table 1) (125 bp PE samples: 21,31,51,71,81,101; 150 bp PE samples 31,51,71,81,101,127; 201FG217 [125 bp PE]: 21,31,51,71). Mitochondrial genomes (mtDNA) were assembled using Novoplasty version 2.7.2 (Dierckxsens et al. 2017), using the Sn15 mitochondrial genome assembly [NCBI RefSeq: <u>EU053989.1</u>] (Hane et al. 2007) as a seed sequence, and with kmers between 31 and 81 (table S5). The k-mer resulting in assemblies with the fewest number of contigs within an expected total assembly size of 47-52 Kb was manually selected and designated the mtDNA sequence of that isolate. Code for generating the mtDNA assemblies is available at https://github.com/darcyabjones/mitoflow.

Nuclear genome assemblies were then filtered for mtDNA sequences by aligning reads to assembled scaffolds using BBmap, and aligning mitochondrial scaffolds to assemblies using minimap2 git commit 371bc95 (Li 2018). Genomic scaffolds were considered to be mitochondrial if the alignment coverage with mitochondrial contigs was greater than 95%, and the median read depth was in the top 0.8% overall read depth. Some manual assignment of very short contigs near the threshold cutoffs was undertaken.

Genome assembly quality control statistics were collected using Quast version 5.0.2 (Gurevich et al. 2013), bbtools stats version 38.38 (https://jgi.doe.gov/data-and-tools/bbtools/bb-tools-user-guide/statistics-guide/), and KAT version 2.4.2 (Mapleson et al. 2017). Code for running post-assembly quality control and selection of mitochondrial scaffolds is available at https://github.com/darcyabjones/postasm (commit: <u>c94c3b9</u>).

### Determining presence-absence variation relative to the SN15 reference isolate

Genome assemblies were aligned to the reference isolate SN15 (Bertazzoni et al. 2021), and to long-read assemblies for isolates SN4 [NCBI Assembly: <u>GCA_002267005.1</u>], SN2000 [NCBI Assembly: <u>GCA_002267045.1</u>], Sn79 [NCBI Assembly: <u>GCA_002267025.1</u>] using nucmer version 4.0.0beta2 (--maxmatch) (Marçais et al. 2018, Richards et al. 2018). Alignments were converted to BED format, from which alignment coverage was computed (bedtools genomecov -bga) and combined (bedtools unionbedg) using BEDTools version 2.28.0 into a bedgraph file (Quinlan and Hall 2010). Mean coverage in non-overlapping 50Kb windows of the genome visualised using the R package circlize (Gu et al. 2014).

Regions of PAV were extracted from the bedgraph using a custom python script (available at: https://github.com/darcyabjones/mumflow/blob/master/bin/find_pavs.py), and coverage blocks were converted to simple presence-absence values (0 or 1) by collapsing coverage count >= 1 to 1. Adjacent blocks with identical PAV values in all isolates were merged, and non-overlapping PAV blocks were also merged where for a set of 3 adjacent blocks, the outer 2 blocks had identical PAV values in all isolates and the centre block was <= 50bp. Code for running these steps is available at https://github.com/darcyabjones/mumflow.

### Annotation of DNA repeats and non-protein coding gene features

Transposable elements (TEs) were predicted using a combination of tools: EAHelitron git commit c4c3dca (https://github.com/dontkme/EAHelitron), LTRharvest (Ellinghaus et al. 2008) and LTRdigest from genometools version 1.5.10 (Steinbiss et al. 2009), MiteFinder git commit 833754b (Hu et al. 2018), RepeatModeler version 1.0.11 (http://www.repeatmasker.org/RepeatModeler), and RepeatMasker version 4.0.9p2 (http://www.repeatmasker.org) using the species “Parastagonospora nodorum”. Putative TE protein coding regions in the genomes were identified using MMSeqs2 version 9-d36de (Steinegger and Söding 2017), searching protein profiles from selected Pfam families (table S6), GyDB families (Llorens et al. 2011), and a custom MSA database based on protein collections in TransposonPSI (http://transposonpsi.sourceforge.net/) and LTR_retriever (Ou and Jiang 2018) (available at: https://github.com/darcyabjones/pante/tree/master/data/proteins).

Predicted TE sequences from EAHelitron, MiteFinder, RepeatModeler, and MMSeqs protein finding were combined and clustered using VSEARCH version 2.14.1 (Rognes et al. 2016) requiring cluster members to have >=70% identity to the cluster seed sequence (-- cluster_fast combined.fasta --id 0.90 --weak_id 0.7 --iddef 0 --qmask dust). Clusters were filtered based on frequency and conservation across the population, requiring presence in >= 4 distinct genomic locations to be considered present in a genome, and requiring the cluster to be present in >= 20% of the total population. Filtered clusters were aligned using DECIPHER version 2.10.0 (Wright 2015), and classified into subtypes using RepeatClassifier (part of RepeatModeler), which was then used as a final customised library to map repeat locations for each assembly with a final round of RepeatMasker. Genes encoding rRNA and tRNA were predicted with RNAmmer version 1.2 (Lagesen et al. 2007) and tRNAscan-SE version 2.0.3 (Lowe and Chan 2016), respectively. These TE and non-coding RNA predictions were used to “soft-mask” genomes using BEDTools (Quinlan and Hall 2010). Code to run these steps is available at https://github.com/darcyabjones/pante/ (commit: <u>2de5d08</u>).

### Annotation of protein-coding genes

Proteins predicted from previous *P. nodorum* SN15 (Bertazzoni et al. 2021), SN4, and SN79 annotations (Richards et al. 2018) were aligned to each genome using Spaln version 2.3.3 (spaln -KP -LS -M3 -O0 -Q7 -ya1 -yX -yL20 -XG20000) (Iwata and Gotoh 2012).

Additionally all fungal proteins from the UniRef 50 database release 2019_08 (https://www.uniprot.org/uniref/, downloaded: 2019-10-29, query: ‘taxonomy:“Fungi [4751]” AND identity:0.5’) were aligned to genomes using Exonerate version 2.4.0 (--querytype protein --targettype dna --model protein2genome --refine region --percent 70 --score 100 --geneseed 250 --bestn 2 --minintron 5 --maxintron 15000 --showtargetgff yes --showalignment no --showvulgar no) (Guy St C and Ewan 2005) with pre-filtering by MMSeqs2 (-e 0.00001 --min-length 10 --comp-bias-corr 1 --split-mode 1 --max- seqs 50 --mask 0 --orf-start-mode 1) (Steinegger and Söding 2017). Published RNAseq data from *P. nodorum* SN15 *in vitro* and 3 days post infection of wheat leaves (Jones et al. 2019) (available in NCBI GEO project: GSE150493; NCBI SRA accessions: SRX8337785, SRX8337784, SRX8337783, SRX8337782, SRX8337777, SRX8337776, SRX8337775, and SRX8337774) were assembled using Trinity v2.8.4 (--jaccard_clip --SS_lib_type FR) (Grabherr et al. 2011) and aligned to genomes using Spaln version 2.3.3 (-LS -O0 -Q7 -S3 - yX -ya1 -Tphaenodo -yS -XG 20000 -yL20) (Iwata and Gotoh 2012), and GMAP version 2019-05-12 (Wu and Watanabe 2005). RNAseq reads were also aligned to all genomes using STAR version 2.7.0e (Dobin et al. 2013) and assembled into transcript annotations using StringTie version 1.3.6 (--fr -m 150) (Pertea et al. 2015).

Genes were initially predicted for each genome using multiple tools: PASA2 version 2.3.3 (-T --MAX_INTRON_LENGTH 15000 --ALIGNERS blat --transcibed_is_aligned_oriented --TRANSDECODER --stringent_alignment_overlap 30.0) (Haas et al. 2003), GeneMark-ET (--soft_mask 100 --fungus) (Lomsadze et al. 2014), CodingQuarry version 2.0 (including the unpublished “Pathogen Mode” with signal peptide predicted using SignalP version 5.0b (Testa et al. 2015, Armenteros et al. 2019), Augustus git commit 8b1b14a (independently for both forward and backward strands; --hintsFile=hints.gff3 --strand=$(Dalman et al.) --allow_hinted_splicesites=’gtag,gcag,atac,ctac’ --softmasking=on -- alternatives-from-evidence=true --min_intron_len=5) (Stanke et al. 2008), and GeMoMa version 1.6.1 (transferring SN15 annotations only) (Keilwagen et al. 2018). Gene predictions using PASA2 used hints from assembled RNASeq transcripts aligned to the genomes with GMAP and BLAT. Augustus gene predictions used transcript alignments by GMAP, intron locations from STAR read alignments, and Spaln and protein alignments by Spaln as hints.

Pan-genomic gene sets may be prone to annotation errors in which orthologous loci are incorrectly annotated in some isolates, leading to false absences. To improve annotation consistency between isolates, protein predictions from PASA, Augustus, and CodingQuarry from all isolates were clustered using MMSeqs2 (90% identity and 98% reciprocal coverage), and annotations corresponding to proteins from representative members of the clusters were transferred to all isolates using GeMoMa as described earlier. Annotations and alignments from Genemark-ET, CodingQuarry, Augustus, PASA, both GeMoMa configurations, Exonerate, Spaln protein and transcript alignments, and GMAP alignments were combined using EVidenceModeler (git commit 73350ce) (--min_intron_len 5) (Haas et al. 2008). Because EVidenceModeler does not support prediction of non-standard splice sites or overlapping genes in different strands, Augustus (with all hints and the same parameters described earlier) was used to predict additional genes in regions of the genomes with hints that didn’t overlap the EvidenceModeler predicted genes on the same strand. Protein predictions were searched against AntiFam (Eberhardt et al. 2012) using HMMER version 3.2.1 (--cut_ga) and matches to pseudogenes were removed. Genes within the merged Augustus and EVidenceModeler gene sets were marked as “low confidence” if supported only by Spaln or GMAP transcript alignments, Exonerate protein alignments, or transfers of annotations between isolates performed via GeMoMa (excluding the initial set of genes transferred from curated SN15 annotations). In SN15, genes only supported by the above tools or Augustus were also marked as low-confidence. “Low-confidence” genes that overlapped other genes on either strand by more than 30% of their length were removed. We corrected errors in the CDS coordinates where phases of gene annotations may lead to incorrect translations in some downstream pipelines by searching against all proteins from other isolates without stop codons, and all Pezizomycota proteins from UniRef-90 (filter: ‘taxonomy: “Pezizomycotina [147538]” AND identity:0.9’; downloaded: 2020-05-13) using blastx version 2.10.0 (-strand plus - max_intron_length 300 -evalue 1e-5) (Camacho et al. 2009). Genes with an in-phase BLAST match lacking internal stop codons were fixed and retained, genes with an out-of-phase BLAST match with internal stop codons were marked as pseudogenes, and those with no BLAST match and internal stops were discarded. Genes overlapping predicted rRNA genes by more than 50% of their length were also discarded, and genes with exons overlapping assembly gaps were split into multiple fragmented genes, where each fragmented annotation was >= 60 bp.

Gene prediction completeness was evaluated for each isolate using BUSCO version 3 (git commit 1554283) using the “pezizomycotina_odb9” dataset (Waterhouse et al. 2018), and additional statistics were collected by genometools version 1.5.10 (Gremme et al. 2013). The updated SN15 annotations were compared to previously published gene annotation versions (Bertazzoni et al. 2021) using ParsEval/AEGeAn version 0.15.0 (Standage and Brendel 2012). The new SN15 annotations were identified using BEDTools, by excluding all new mRNA predictions overlapping original SN15 “A” and “B” mRNA annotations from (Bertazzoni et al. 2021) on the same strand >= 20% by length (bedtools subtract -a new -b old -s -A -F 0.2), and were designated as the “C” gene.

### Orthology & positive selection

Orthology relationships for predicted proteins were predicted using Proteinortho version 6.0.30 (-singles -seflblast) (Lechner et al. 2011) with using Diamond version 2.0.8 (Buchfink et al. 2021). Alternative isoforms present in the SN15 annotations were included in the orthology finding. Orthologous clusters were assigned identifiers prefixed by “SNOO’’ (supplementary table S10), where clusters with members in isolate SN15 were assigned numbering corresponded to “SNOG” locus numbers (Bertazzoni et al. 2021) (Complete data: https://doi.org/10.6084/m9.figshare.12966971.v3). SNOO groups were also assigned alphabetical prefixes where one or more isoforms were present in SN15. For example “SNOO_434350AB” contains both *SNOG_434350* isoforms A and B, and “SNOO_033200A149040A” contains both SNOG_033200A and SNOG_149040A. Clusters without members in SN15 were assigned sequential numbers starting from 50,000. For the purpose of analysis of selection pressure, a representative SN15 isoform was selected for each SNOO orthogroup by selecting the isoform with the closest length to the mean sequence length of the orthogroup. Representative CDS sequences of SNOO orthogroups were codon-aligned using DECIPHER version 2.16.1 (Wright 2015), and gene trees were estimated using FastTree version 2.1.11 (Price et al. 2010). The SNOO orthogroup codon multiple sequence alignments and the gene trees were used to test for positive selection in the orthogroups using the BUSTED method in the HYPHY package version 2.5.15 (Pond et al. 2005, Murrell et al. 2015). A p-value threshold of 0.01 was used to determine positively selected SNOO orthogroups. Position specific positive selection tests were performed for the known effectors ToxA, Tox1, and Tox3 using the FUBAR method in the HYPHY package (Pond et al. 2005, Murrell et al. 2013).

For the purposes of PAV and pan-genomic comparisons, to account for alternate isoforms present across multiple orthogroups, temporary “locus groups” were constructed by combining orthogroups that share common loci. Copy numbers of locus groups were calculated for each isolate as the number of distinct loci. Large PAV regions were identified by heriarchically clustering locus groups and samples by copy number using UPGMA clustering and the manhattan distance metric for orthogroups and 1 - Pearson’s correlation coefficient as a distance metric for isolates. Manually-selected clusters of locus groups showing PAV were aligned to the SN15 genome where possible using MashMap v2.0 (-s 500 –filter_mode none) (Jain et al. 2018). Orthogroups were designated as “core” if all isolates contained at least one member in the parent locus group, as “accessory” if more than one isolate but not all isolates had at least one member in the locus group, and as “singleton” if the locus group was only detected in a single isolate. Additionally, locus groups were designated as “multicopy” if any isolate had more than one member.

### Functional analysis & effector candidate prediction

Predicted whole protein functions were found by searching the Swiss-Prot database version 2020_02 (Bairoch and Apweiler 2000) using MMSeqs2 version 11-e1a1c (--start-sens 3 -s7.5 --sens-steps 3 -a) (Steinegger and Söding 2017). Matches were considered reliable for functional annotation if they covered >= 70% of both sequences, with >=60% sequence identity, and an e-value < 1e-10. Functional domains were predicted using InterProScan (Jones et al. 2014, Mitchell et al. 2019). Additionally, GO-terms and predicted product names were predicted using the web-servers of PANNZER (Koskinen et al. 2015) and eggNOG-Mapper (Huerta-Cepas et al. 2017). GO-term predictions from InterProScan, PANNZER, and eggNOG-Mapper were combined and filtered to exclude terms in the GO do_not_annotate “anti-slim” set (available at: http://geneontology.org/docs/download-ontology, downloaded: 2020-05-15) to remove uninformative terms, forming the final GO-term set for the predicted proteomes.

Effector-like sequences were predicted using the Predector pipeline (https://github.com/ccdmb/predector, version: 0.1.0-alpha), which incorporates several software analyses including SignalP versions 3.0, 4.1g, 5.0b (Armenteros et al. 2019), DeepSig (Savojardo et al. 2018), TargetP version 2.0 (Armenteros et al. 2019), DeepLoc version 1.0 (Almagro Armenteros et al. 2017), TMHMM version 2.0c (Krogh et al. 2001), Phobius version 1.01 (Käll et al. 2004), EffectorP versions 1 and 2 (Sperschneider et al. 2016, Sperschneider et al. 2018), ApoplastP version 1 (Sperschneider et al. 2018), LOCALIZER (Sperschneider et al. 2017), homology searches against dbCAN version 8 using HMMER version 3.3 (Yin et al. 2012, Mistry et al. 2013), and sequence matches against PHI-base version 4.9 (Urban et al. 2020) using MMSeqs2 version 11.e1a1c (Steinegger and Söding 2017). Information from Predector, InterProScan, Pannzer, eggNOG-mapper, positive selection and orthogroup analyses were combined into a single table (summarised in table S10. Complete data available at https://doi.org/10.6084/m9.figshare.12966971.v3).

Enriched and depleted GO terms were detected for core, accessory, and singleton orthogroups (and their multicopy subsets), accessory orthogroups contained in between 20% and 80% of isolates, orthogroups predicted to be under positive selection anywhere in the gene tree and with more than 20% of members, for orthogroups not found in SN15, manually selected clusters of orthogroups identified from hierarchical clustering (Table S9), *P. nodorum* Sn15 genes on accessory chromosome 23, and *P. nodorum* Sn15 genes with a ratio of RIP-like mutations over transitions within 1000 bp of the gene over the 95th percentile, using two-tailed hypergeometric tests implemented in the GOATOOLS version 1.0.6 (Klopfenstein et al. 2018). For each of these sets, two-tailed hypergeometric/Fisher’s exact tests were also used to test for enrichment of genes lacking any GO term assignments, genes annotated as secreted by the Predector pipeline, and genes annotated as secreted and with a positive EffectorP 2 prediction using the SciPy Python package (Virtanen et al. 2020).

### Data availability

All sequencing data, genomes and annotations generated are available under NCBI Bioproject: PRJNA612761. Gene annotations for isolates SN2000, SN4, and SN79, as well as short variant predictions relative to SN15 are deposited online at (https://doi.org/10.6084/m9.figshare.13340975). Complete functional annotations, orthogroup assignments, CDS alignments and trees, and positive selection tests are available online at (https://doi.org/10.6084/m9.figshare.12966971.v3}. Supplementary material is also available online at (https://doi.org/10.6084/m9.figshare.13325915.v2).

## Author Statements

### Authors and Contributors

DABJ, JKH, K-CT, and HTTP conceived and designed the study. KR and HTTP performed laboratory work. DABJ, SB, and JKH performed bioinformatics and data analysis. DABJ and JKH wrote the manuscript. DABJ, K-CT, HTTP, and JKH edited the manuscript. All authors read and approved the manuscript.

### Conflicts of Interest

The authors declare there are no conflicts of interest.

## Funding Information

This study was supported by the Centre for Crop and Disease Management, a joint initiative of Curtin University and the Grains Research and Development Corporation (Research Grant CUR00023). This research was undertaken with the assistance of resources and services from the Pawsey Supercomputing Centre and the National Computational Infrastructure (NCI), which is supported by the Australian Government. This research is supported by an Australian Government Research Training Program (RTP) Scholarship.

### Ethical Approval

n/a

